# Unique dental arrangement in a new species of *Groenlandaspis* (Placodermi, Arthrodire) from the Middle Devonian of Mount Howitt, Victoria, Australia

**DOI:** 10.1101/2024.09.11.612576

**Authors:** Austin N. Fitzpatrick, Alice M. Clement, John A. Long

**Affiliations:** College of Science and Engineering, Flinders University, Adelaide, South Australia, Australia

## Abstract

Well-preserved specimens of an undescribed species of arthrodiran placoderm, *Groenlandaspis howittensis* sp. nov. (Middle Devonian of Victoria, Australia), reveals previously unknown information on the dermal skeleton, body-shape and tooth arcade of the wide-spread genus *Groenlandaspis*. The new material includes, dual pineal plates, extrascapular plates, and cheek bones cheek bones showing the presence of cutaneous sensory organs. The anterior supragnathal, usually a paired element in arthrodires, is a fused medial bone in *G. howittensis* sp. nov. It is positioned anterior to the occlusion of the mouth between the lower jaw (infragnathals) and upper jaw (posterior supragnathals) bones, indicating a specialised feeding mechanism and broadening the known diversity of placoderm dental morphologies. *G. howittensis* sp. nov. differs from all other groenlandaspidids by a less pronounced posterior expansion of the nuchal plate; the shape of the posterior dorsolateral plate and the presence of a short accessory canal on the anterior dorsolateral plate. A new phylogenetic analysis positions Groenlandaspididae in a monophyly with the phlyctaeniid families Arctolepidae and Arctaspdidae, however, the specific intrarelationships of groenlandaspidids remain poorly resolved.

## INTRODUCTION

Arthrodires are an extinct clade of placoderms (stem-jawed vertebrates) and a dominant faunal component of Devonian marine and freshwater ecosystems. Arthrodires are one of the earliest jawed vertebrates to show evidence of true teeth (Smith & Johanson 2003; Rücklin et *al.* 2012; Vaškaninová *et al*. 2020) and provide valuable insight into the early evolution of feeding ecologies, including durophagy (Dennis & Miles 1979), suspension feeding (Coatham *et al*. 2020) and pelagic hunting strategies (Jobbins *et al*. 2024). However, knowledge of these specialisations is generally limited to more derived forms, such as the Eubrachythoraci, which possess more robust jaw bones. Consequently, the morphology of more basal forms, such as that of the globally occurring family Groenlandaspididae, remain poorly understood.

Groenlandaspidids are known from Lower to Upper Devonian deposits throughout Gondwana (Young 1993; Anderson *et al*. 1999), attaining a cosmopolitan distribution following a northward dispersal into Laurussia in the Late Devonian (Janvier & Clément 2005). The namesake genus, *Groenlandaspis*, Heintz 1932, is the most diverse consisting of 10 named species (Heintz, 1932; Ritchie, 1975; Janvier & Ritchie, 1977; Chaloner *et al*., 1980; Long *et al*. 1997; Daeschler, Frumes & Mullison 2003; Janvier & Clément, 2005; Olive *et al*, 2015) and numerous more occurrences categorised only to genus level (Young 1993).

The Middle Devonian Mount Howitt, fossil site (Victoria, Australia) preserves a diverse freshwater fish fauna (Table. 1) as compressed articulated individuals displaying aspects of both dermal and visceral morphology (Long, 1983a; 1983b; 1984; 1986a; 1986b; 1987; 1988; 1992; 1999; Long & Holland, 2008; Long & Clement 2009; Holland, Long & Snitting, 2010). We herein describe well-preserved and extensive material of a new species, *Groenlandaspis howittensis* sp. nov., representing the first member of the globally-distributed family to be formally described from Australia. This new material reveals undescribed features of the tooth plates, squamation and body-shape of the genus.

Multiple characteristics have been suggested to be important for the evolution of groenlandaspidids (Long 1995; Olive *et al*. 2015) but none have been incorporated into a computer driven analysis until now.This new complete material such as this offers the opportunity to clarify the phylogenetic relationships of *Groenlandaspis,* and the intra and interrelationships of Groenlandaspididae. The phylogenetic relationships of Devonian fish have been used to infer the geographic dispersal patterns of vertebrate groups, as has been recently demonstrated for bothriolepid antiarch placoderms (Dupret *et al*. 2023).

## MATERIALS AND METHODS

Fossil preparation — Specimens were collected from Taungurong country, Victoria, during field trips lead by Professor Jim Warren of Monash University between 1970-1974, and by the late Alex Ritchie of the Australian Museum in the early 1990’s. The *Groenlandaspis* material consists of specimens from the upper conglomerate and lower mudstone units of the Bindaree Formation (Long, 1983a). Specimens were prepared in 15% Hydrochloric acid (HCl) solution to dissolve friable bone to reveal both sides preserved of an individual as impressions within the rock. Black latex casts were whitened with ammonium chloride to reveal fine anatomical detail for comparative analysis.

### Phylogenetic analysis

To investigate the evolutionary relationships of the genus *Groenlandaspis* and the family Groenlandaspididae we performed a phylogenetic analysis of selected phlyctaenoid arthrodires using a morphological character matrix modified from the matrix of 121 characters and 60 taxa of Zhu *et al*. (2016). 11 new characters were identified from the literature or during the course of this research and incorporated in this existing matrix (Table 2), forming a new matrix of 132 characters and 72 taxa. The matrix was treated with MESQUITE v3.61 (Maddison & Maddison 2019), some corrections were made (see appendix). In addition to *G. howittensis* sp. nov. described herein, nine more taxa were added to the ingroup, including the type species for *Groenlandaspis*, *G. mirabilis*, Heintz 1932 and four relatively complete groenlandaspidids: *Tiaraspis subtilis*, (Gross, 1933), *Groenlandaspis riniensis* and *Africanaspis doryssa,* Long *et al*., 1997, and *Mulgaspis evansorum*, Ritchie, 2004. As well as two arctolepidids (*Arctolepis decipiens*, (Woodward, 1891), and *Heintzosteus brevis*, (Heintz, 1929)). Two selenosteids, *Alienacanthus malkowskii*, Kulczycki, 1957 and *Amazichthys trinajsticae*, Jobbins *et al*. 2022, were added for diversity.

**Table. 1:**
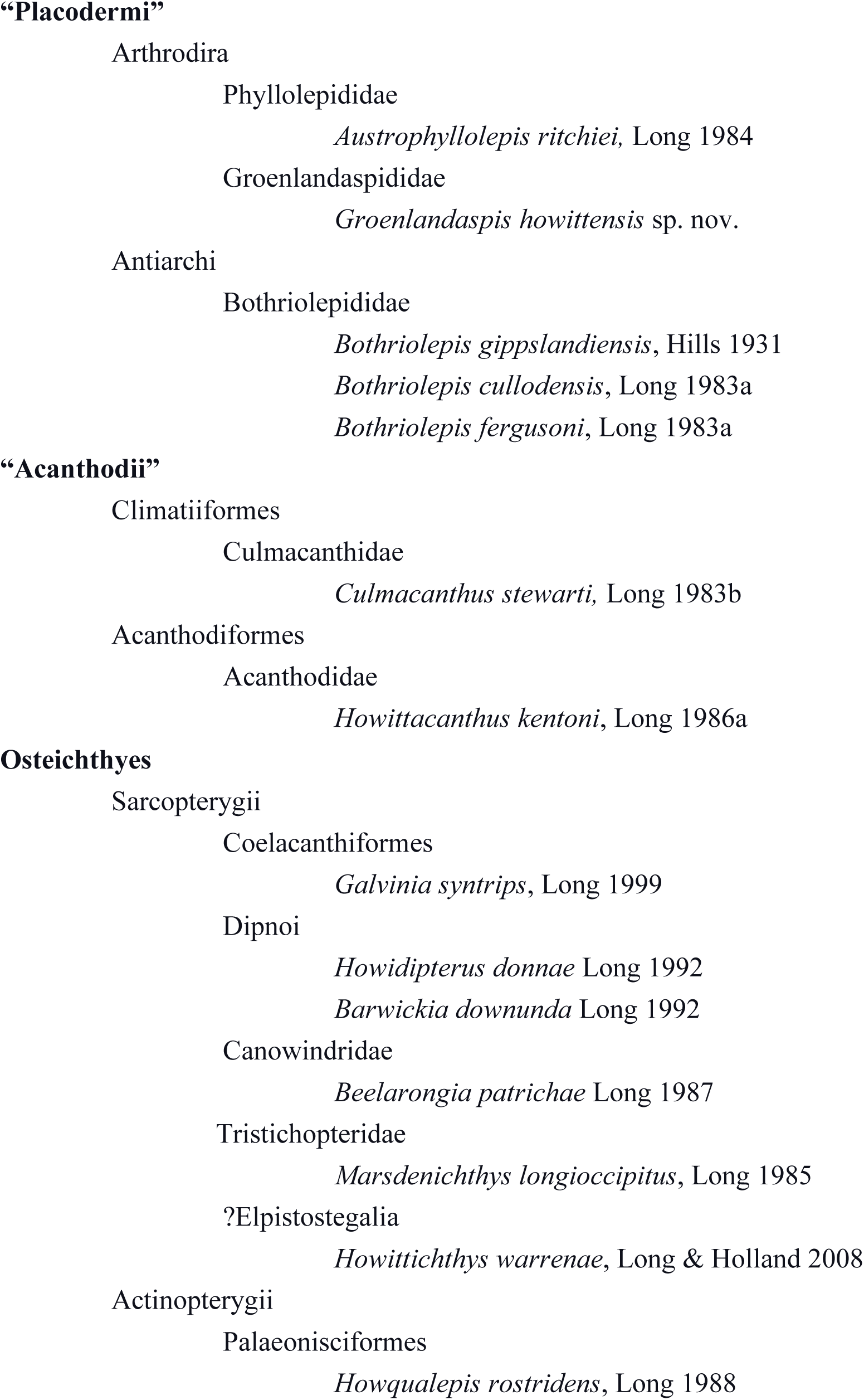
Faunal List from the Mount Howitt locality, Victoria, Australia following Long 1983; 1999.

**Table 2.** 11 new characters added onto a matrix of 121 characters from Zhu *et al*. (2016).

| *No.* | *Description* | *Reference* |
| --- | --- | --- |
| 122 | Cervical Joint: Sliding (0) Ginglymoid (1). | Miles 1973 |
| 123 | Transversely divided pineal plate forming anterior and posterior plates: Absent (0) Present (1). | This article |
| 124 | Cutaneous sensory pits present on the suborbital or/and post suborbital plates: Absent (0) Present (1). | King, Hu & Long, 2016 |
| 125 | Dermal contact between the anterior dorsolateral and posterior lateral plates: Absent (0) Present (1). | This article |
| 126 | Inverted V-shaped flexure of the posterior dorsolateral plate sensory canal. Scored not applicable in taxa without a PDL sensory canal: No flexure (0) Weak flexure, >110° (1) Strong flexure, <110° (2). | Long 1995 |
| 127 | Dorsolateral ridge originating from near the condyle of the anterior dorsolateral plate: Absent (0) Present (1). | Long 1995 |
| 128 | Medial contact of the dorsolateral plates under the median dorsal plate: No contact (0) anterior dorsolateral plates (1) anterior and posterior dorsolateral plates (2). | Goujet 1984 |
| 129 | Internal annular thickening of the posterior trunk plates ('b.cpd', Goujet 1984, fig. 61B): Absent (0) Present (1). | Goujet 1984 |
| 130 | Median contact of the posterior ventrolateral plate: Simple overlap (0) Sigmoidal/double overlapping (1) | Goujet 1984, Dupret 2004 |
| 131 | Ventral sensory canals: Absent (0) Present (1) | This article |
| 132 | Distinct infraspinal lamina/process ('pr.infsp', Miles & Westoll 1968, fig. 40C; 'la.spv', Goujet 1984, fig. 66A) of the anterior ventrolateral plate: Absent (0) Present (1). | This article |
| 133 | Anterior ventral plates: Absent (0) Present (1) | Miles 1973 |

Using our modified matrix, a phylogenetic analysis was performed in PAUP* 4.0 (Swofford, 2003) using a heuristic search with a random addition sequence of 1000 repetitions and holding 1000 trees per search. Characters 4, 14, 20, 35, 51, 75, 92, 93, 126, and 128 were ordered as they form a morphoclines. The tree was rooted using the actinolepid arthrodires *Kujdanowniaspis podolica*, (retained from Zhu *et al*. (2016)) and two additional taxa, *Lehmanosteus hyperboreus*, Goujet, 1984, and the genus *Bryantolepis*, scored as a composite of the species *Bryantolepis brachycephela*, Camp,*et al*. 1949, and *Bryantolepis williamsi*, Elliot & Carr, 2011. Outgroup taxa were selected for their completeness and sister relationship to Phlyctaenoidei, see the phylogenetic analyses of Dupret (2004) and Dupret *et al*. (2017).

### Amended Diagnosis

Groenlandaspidids with pineal element either singular or divided into dual anterior and posterior plates (APi and PPi); rostrally developed preorbital plates that contact the suborbital plate; postnasal plates absent. Extrascapular plates overlying a shallow posterior descending lamina. Dorsoventrally flattened upper tooth-plates consisting of a fused, crescentric, anterior supragnathal and paired posterior supragnathals. Anterior ventral plates absent. Large posterior dorsolateral plate with sharp V-shaped flexure of the lateral canal (<110◦). Median dorsal plate longer than high.

### Remarks

The generic diagnosis has not been updated since Stensiö, (1939) described material of *Groenlandaspis* from East Greenland, then only consisting of the type species, *G. mirabilis*. Thereafter, additional species have been referred to the genus based on general resemblance, and researchers have since suggested that the genus does not represent monophyletic clade (Janvier & Clément, 2005; Olive *et al*., 2015). *GROENLANDASPIS HOWITTENSIS* sp. nov.

### Diagnosis

Medium sized *Groenlandaspis* with an adult armour length up to 150mm and a reconstructed total body length of approximately 300mm. Skull-roof as long as broad with gently concaved posterior margin. Anterior dorsolateral plate possessing a short dorsal accessory canal. Posterior dorsolateral plate higher than long (NMV P48875, H/L = 1.44); lateral canal sharply flexed (between 96°, NMV P48875 and 105°, AMF 62437). Median dorsal plate sub-equilateral (H/L = approx. 0.65); caudal margin gently concaved and lined with prominent tubercules.

### Etymology

After the site where it was found at the base of Mount Howitt

### Holotype

NMV P48873, a complete specimen showing a flattened and complete headshield with partial lateral trunk shield and pectoral fin preserved (Fig. 1A, C).

**Figure 1.**
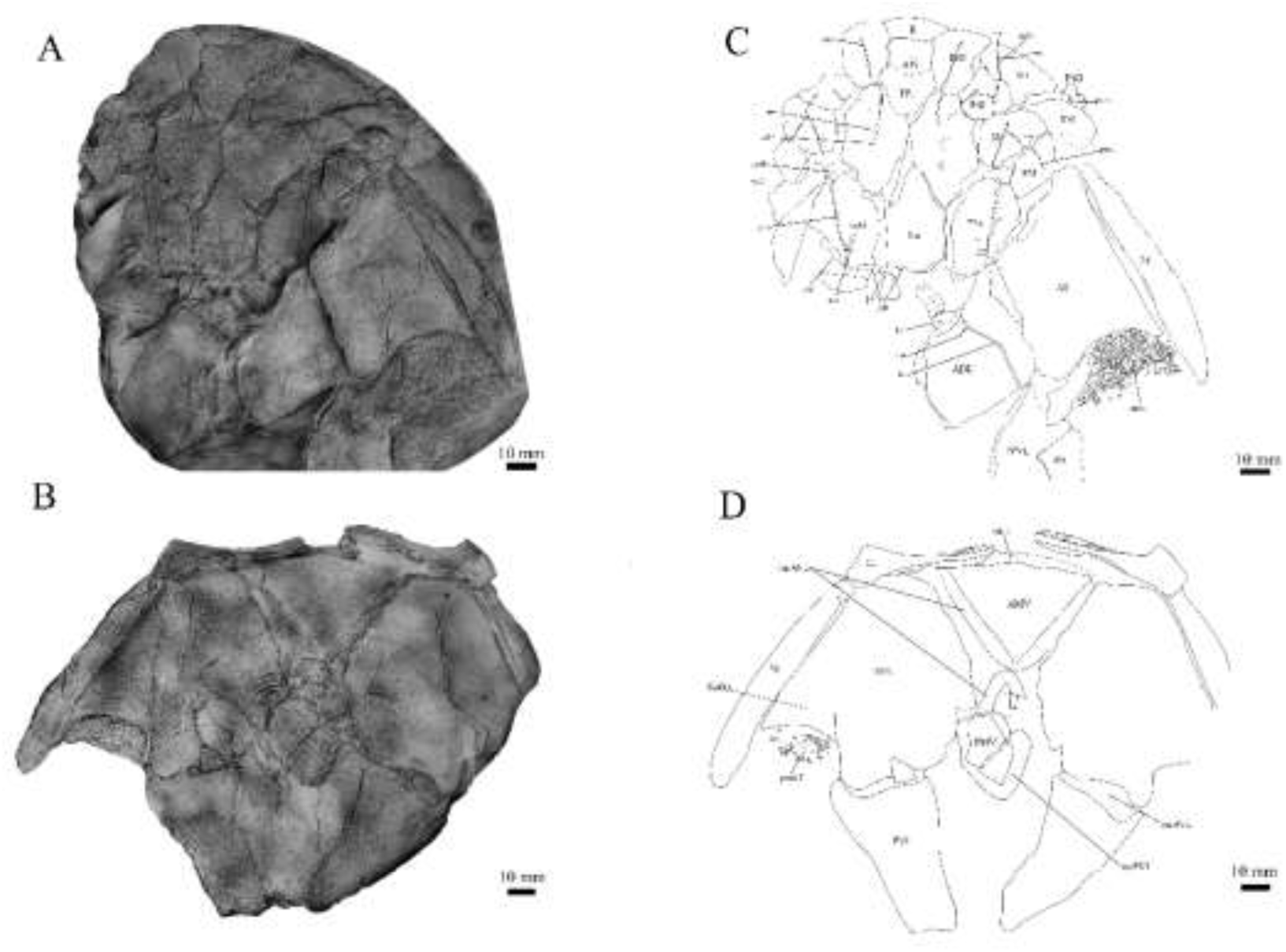
*G. howittensis* sp. nov. photos A, the holotype NMV P48873, head shield and partial trunk shield in dorsal view; B, trunk shield in ventral view, NMV P48874. Latex peels whitened with ammonium chloride. C, D, sketch interpretations of same specimens.

### Referred Specimens

NMV P48874, counterpart to the holotype showing a complete ventral trunk shield (Fig. 1B, D) and tooth plates (Fig. 2) preserved in life position.

**Figure 2.**
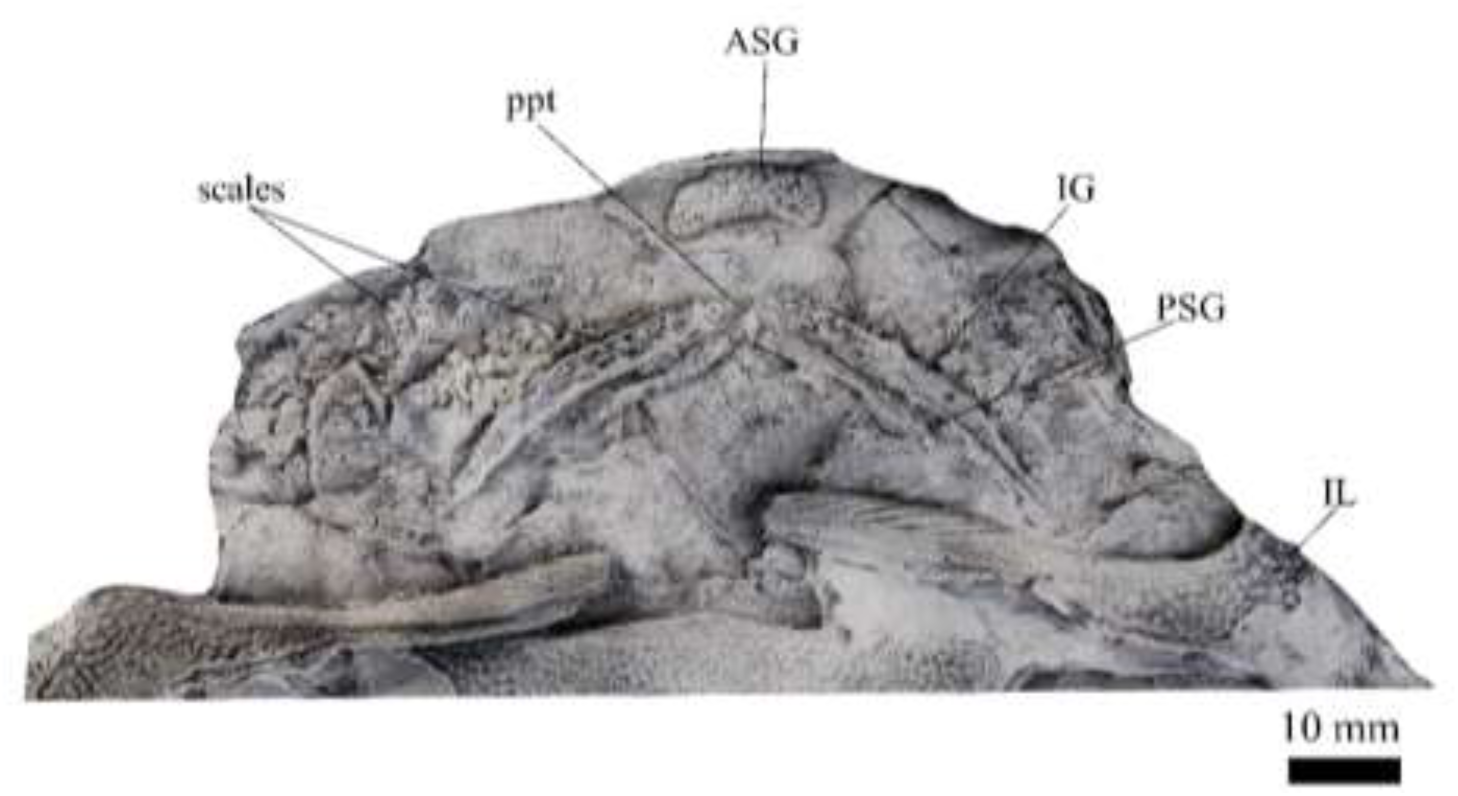
*G. howittensis* sp. nov., jaws of NMV P48773 in ventral view. Latex peel whitened with ammonium chloride.

### Locality, Horizon, and Age

*G. howittensis* sp. nov. remains are known from the upper sandstone conglomerate and lower mudstone shale members of the Bindaree Formation exposed at the Mount Howitt Spur fossil site (Long 1983a). The holotype derives from the lower shale member. The age of the Mount Howitt fauna is considered to be Givetian based on evidence of its faunal composition and comparison with other Devonian fish faunas in south-eastern Australia (Young, 1993; 2007; Long, 1999; Long *et al*., 2021).

## RESULTS

### Description

#### Skull roof

The skull roof of *G. howittensis* sp. nov. is known from several complete and partial specimens (Fig. 1, 3, 5, 6). It is overall very similar to *G. antarcticus* (Ritchie, 1975) but differs by its more deeply situated orbits and nuchal plate. The cranial sensory canals adhere to the pattern described in other species of *Groenlandaspis* where complete crania are known, *G. antarcticus* and *G. riniensis* (Ritchie 1975; Long *et al*., 1997). Other species of *Groenlandaspis* show no evidence of post nasal bones and we suspect they are completely reduced as in *Arctolepis* (Goujet, 1984). The pineal element of *G. howittensis* sp. nov. is formed of anterior (APi) and posterior pineal (PPi) plates, and in articulation they form approximately one third of the cranial length (Fig. 1A, C). In the holotype of *G. howittensis* sp. nov. the APi and PPi are fused and the suture is faint but several other specimens clearly show both plates in association but disarticulated (Fig. 4).

**Figure 3.**
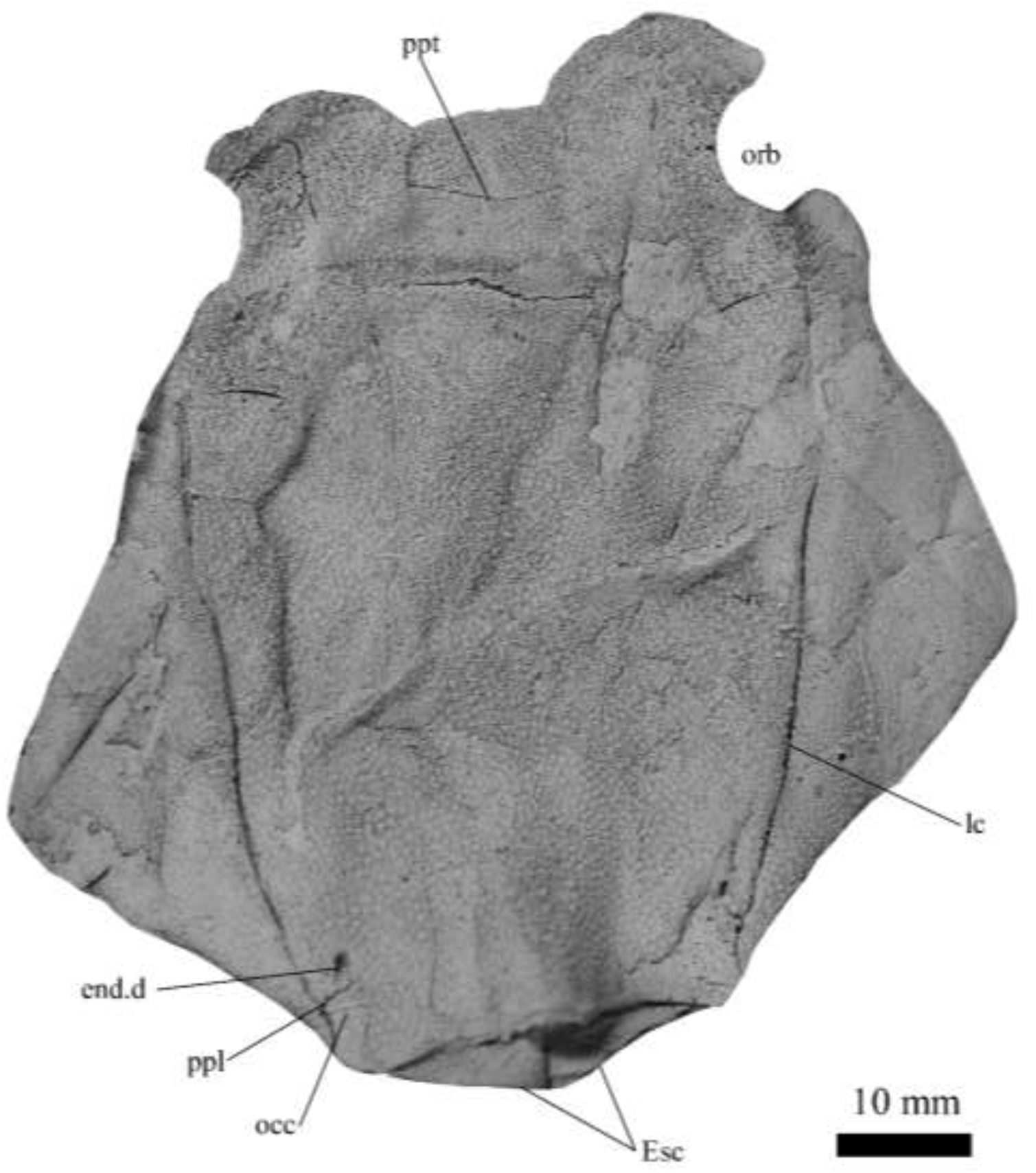
*G. howittensis* sp. nov., AMF 63548, skull roof in dorsal view. Latex peel whitened with ammonium chloride.

**Figure 4.**
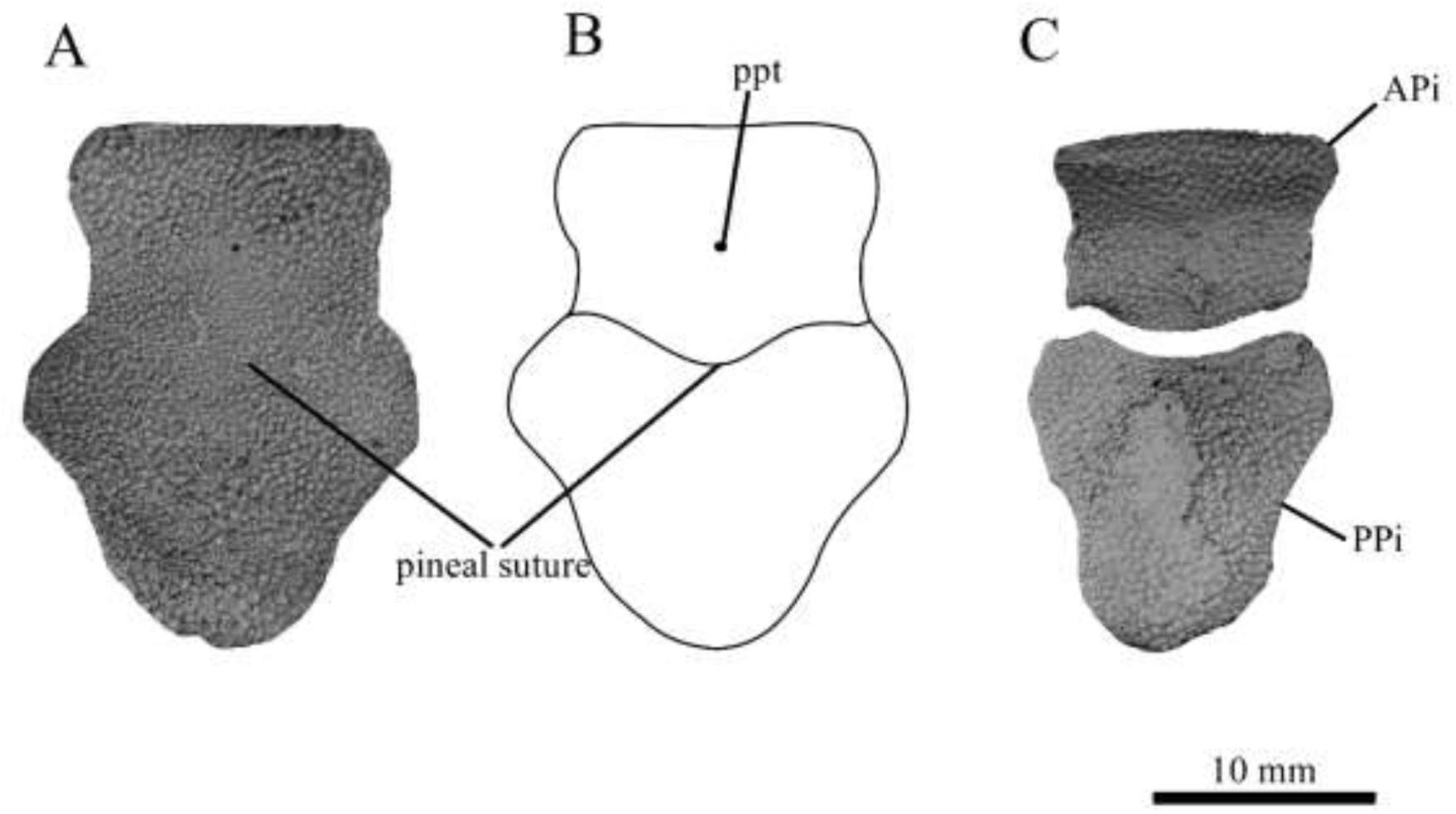
*G. howittensis* sp. nov., pineal plates in dorsal view. A, Pineal plate of NMV P48873, B, interpretive drawing of the same. C, APi and PPi of AMF 62532. A, C, Latex peels whitened with ammonium chloride.

**Figure 5.**
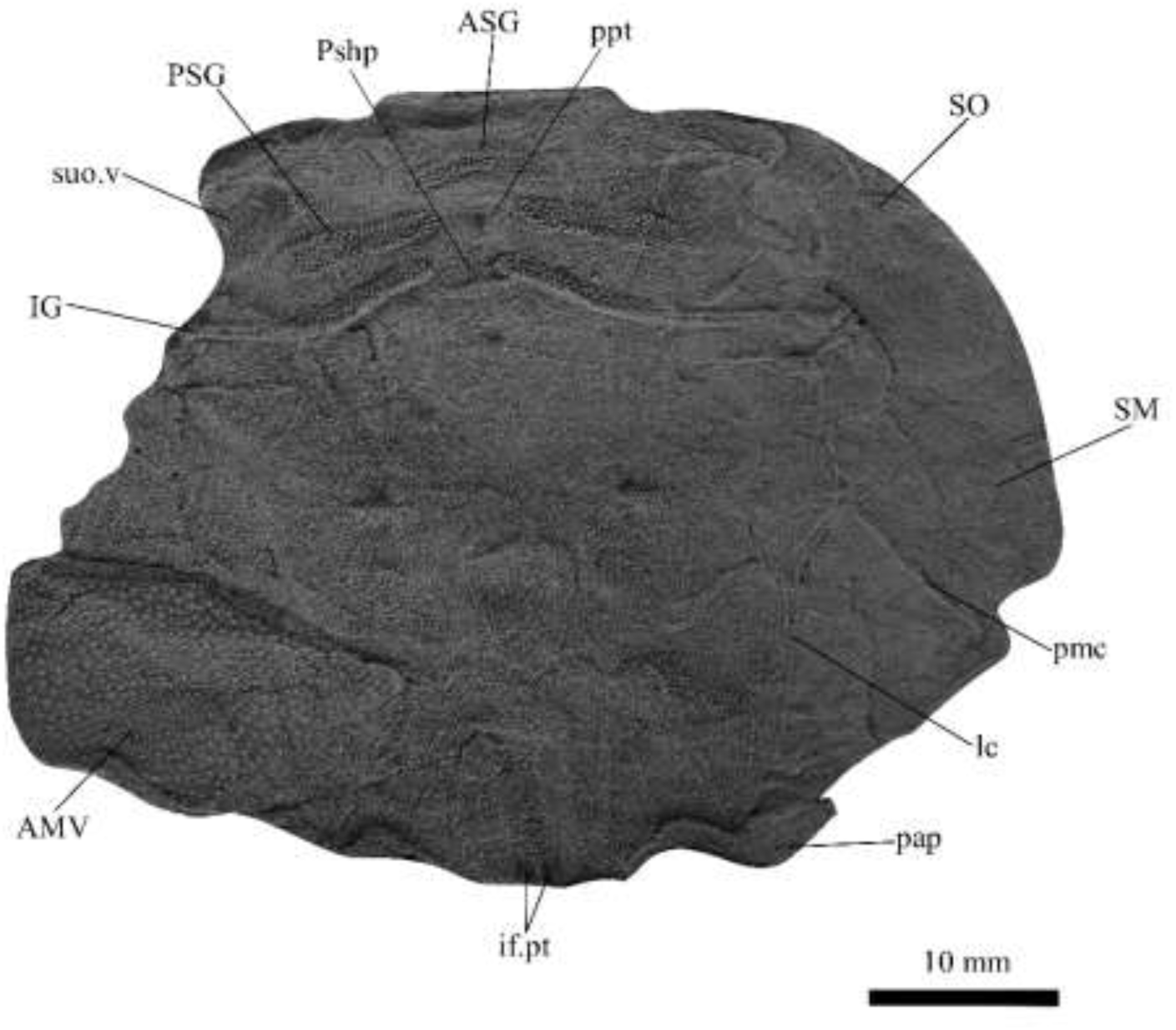
*G. howittensis* sp. nov., AMF 62534, juvenile, head shield in ventral view. Latex peel whitened with ammonium chloride.

**Figure 6.**
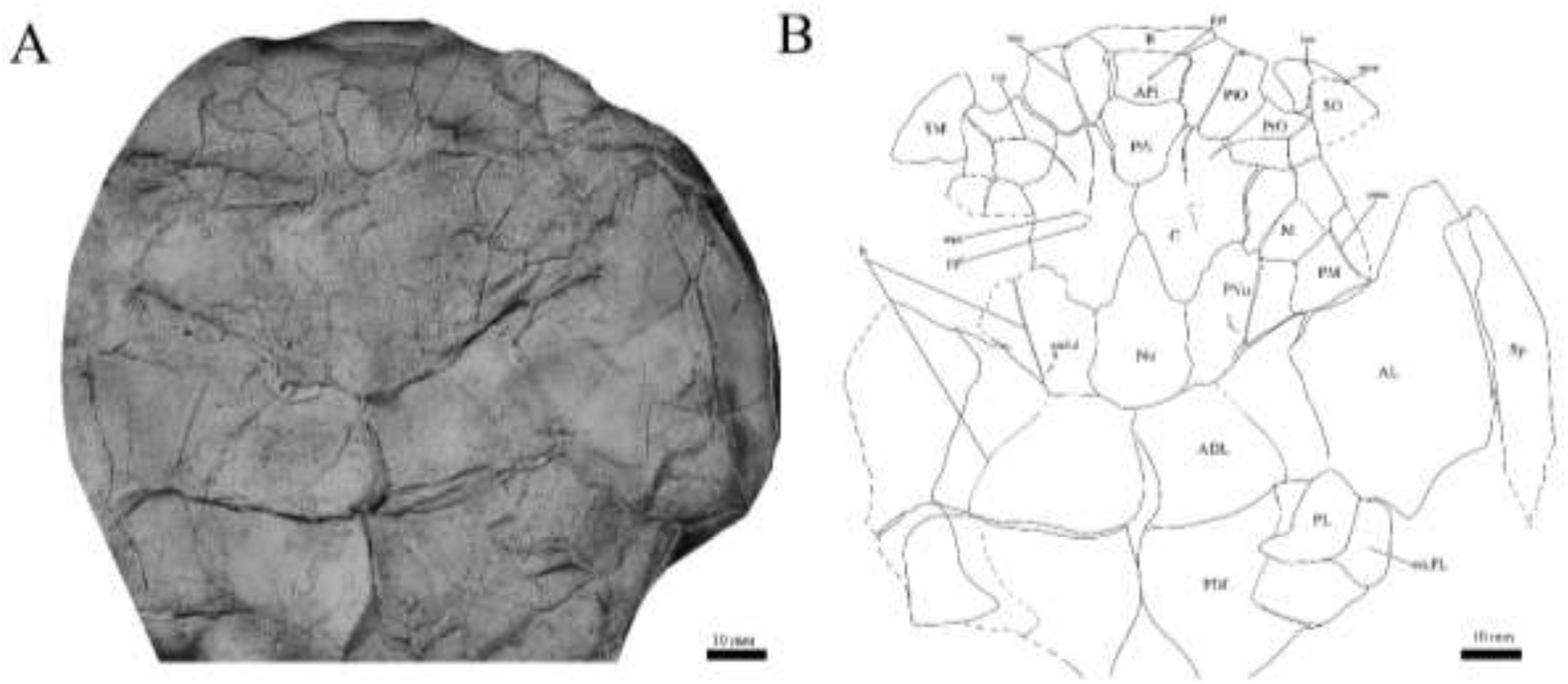
*G. howittensis* sp. nov. head and trunk shield in dorsal view. A, AMF 62532, latex peel whitened with ammonium chloride. B, interpretive line drawing of same specimen, dotted lines indicate broken or incomplete plate margins.

**Figure 7.**
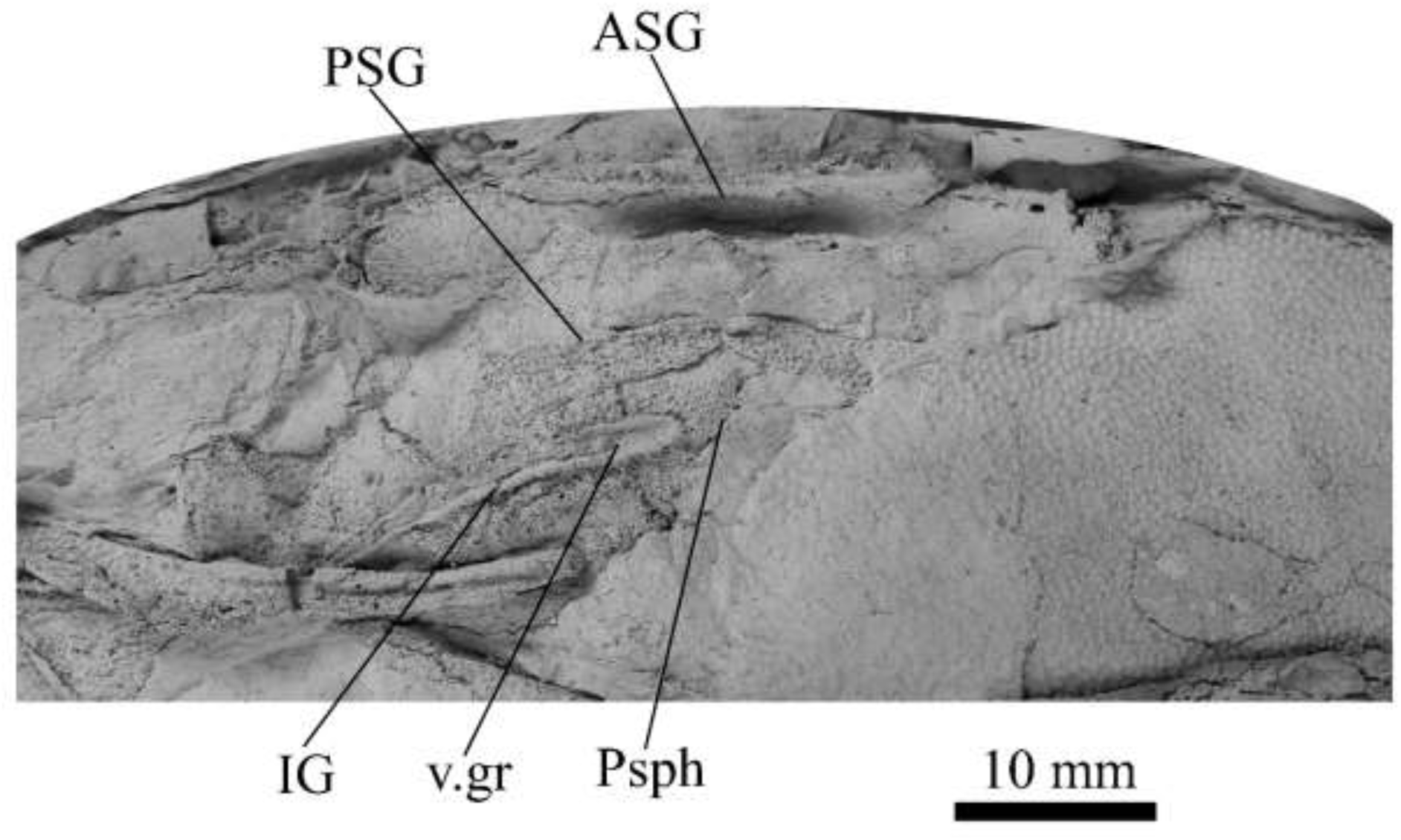
*G. howwitensis* sp. nov. tooth plates in ventral view, AMF 62333, latex peel whitened with ammonium chloride.

Dual pineal plates are a distinct feature in some members of the Groenlandaspididae and, thus far, one or both plates have also been described for *Turrisaspis Africanaspis*, *Colombiaspis* (Olive *et al*., 2015; 2019; Gess & Trinajstic, 2017) and are presumed to be present in *Tiaraspis* based on the gap in the headshield once reconstructed (Schultze, 1984). Dual pineal plates are herein described for the first time in a species of *Groenlandaspis* but have been previously noted in other species: *G. disjectus*, *G. antarcticus* and *Groenlandaspis* sp. from Canowindra, New South Wales, Australia (Ritchie, 2004, and pers. obv.) but are not confirmed for *G. riniensis* from the Waterloo Farm Lagerstatte, South Africa. The central plates are essentially identical to *G. antarcticus* differing only in a further developed embayment area for the postorbital plate (PtO). The nuchal (Nu) plate is longer than broad (B/L = 0.6, NMV 48874, Fig. 1A, C) and is roughly 40% of the cranial length, it is transversely convex, rising posteriorly to a slight median crest. The plates posterior margin is enwrapped by small postnuchal processes of the paranuchal plates (PNu). Extrascapular plates (ESC) are preserved within the nuchal gap of one articulated specimen (Fig. 3) and a fragment of a possible dissociated ESC is also identified in AMF 155378 (Fig. 8). As in brachythoracids, e.g. *Millerosteus minor* (Desmond, 1974, fig. 1C), the extrascapulars are paired plates which overlie the posterior descending lamina (pdl) of the skull-roof (Fig. 2C) and are furrowed by a sensory canal, unlike brachythoracids, this sensory canal does not converge with the occipital cross commissure (occ) of the PNu, instead arcing posteriorly, possibly aligning with the dorsal accessory canal (acc) of the ADL plate. The visceral surface of the skull-roof (Fig. 5, 10) displays no continuous nuchal or occipital thickening as developed in brachythoracids though infranuchal pits (if.pt) are present, as in *Parabuchanosteus* (Young, 1979) and many other taxa.

**Figure 8.**
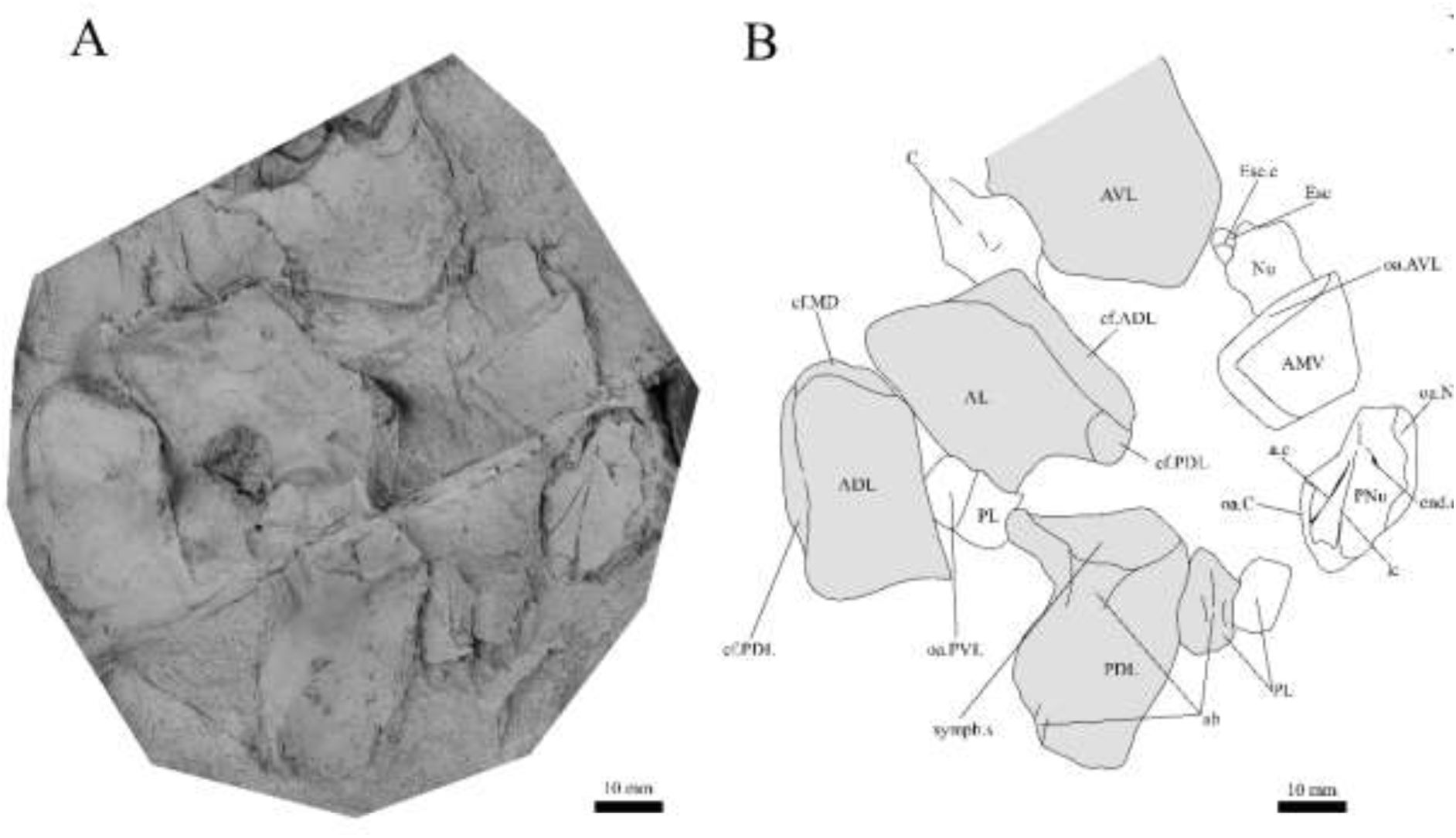
*G. howittensis* sp. nov., disarticulated head and trunk plates in multiple views, A, AMF 155378, latex peels whitened with ammonium chloride. B, interpretive drawing of the same specimen. shaded areas indicate the internal side of the plate.

### Cheek plates

The cheek unit comprises of large submarginal (SM) and suborbital plates (SO) divided by a slender post suborbital plate (PSO). The suborbital lamina of the SO which encloses the ventral portion of the orbit is short and deep and contacts the PrO as in some eubrachythoracids, e.g. *Eastmanosteus* (Dennis-Bryan, 1987). The dermal surface of the plate carries two deep sensory lines, the supraoral (sorc) and infraorbital canals (ioc), which meet in the radiation centre of the plate (Fig. 6). In some individuals, such as in the holotype (Fig. 1), the supraoral canal terminates just before meeting the infraorbital canal into a cutaneous pit (cu.so). The PSO is preserved in the holotype with the ventral portion of the plate broken and disarticulated (Fig. 1). The PSO is a slender bone which tightly situates into the posterior notch of the SO plate, its dermal surface is furrowed longitudinally by postorbital sensory canal (psoc). The submarginal plate (SM) is preserved close to life position but broken in the holotype; in one near complete specimen the SM is complete and displaced anterior to its life position and better reveals its overall shape (Fig. 6B). The SM of *G. howittensis* sp. nov. is the first of example of this bone described for a groenlandaspidid. It is a large, ellipsoidal bone which overlapped the lateral margin of the skull roof and postbranchial lamina of the AL plate, as in other basal arthrodiran forms, e.g. *Wuttagoonaspis* and *Dicksonosteus* (Ritchie, 1973; Goujet, 1984).

### Tooth plates

The tooth plates are preserved as impressions in several specimens (Figs. 2, 5, 7, 10), but are best represented in the counterpart of the holotype where the infragnathals (IG) are superimposed onto the posterior supragnathals (PSG) (Fig. 2). The tooth plates do not exhibit any wear facets as noted for eubrachythoracids like *Dunkleosteus* (Lebedev *et al*., 2023). The crescentic denticulated bone positioned under the rostral plate in this specimen and others is interpreted here as a fused anterior supragnathal (ASG) derived from the ancestral paired condition of other arthrodires, e.g. *Coccosteus* (Miles & Westoll, 1968, fig. 17A,). In one smaller individual the ASG is much slenderer in proportions, suggesting positive allometric growth in this element through ontogeny (Fig. 6).

**Figure 9.**
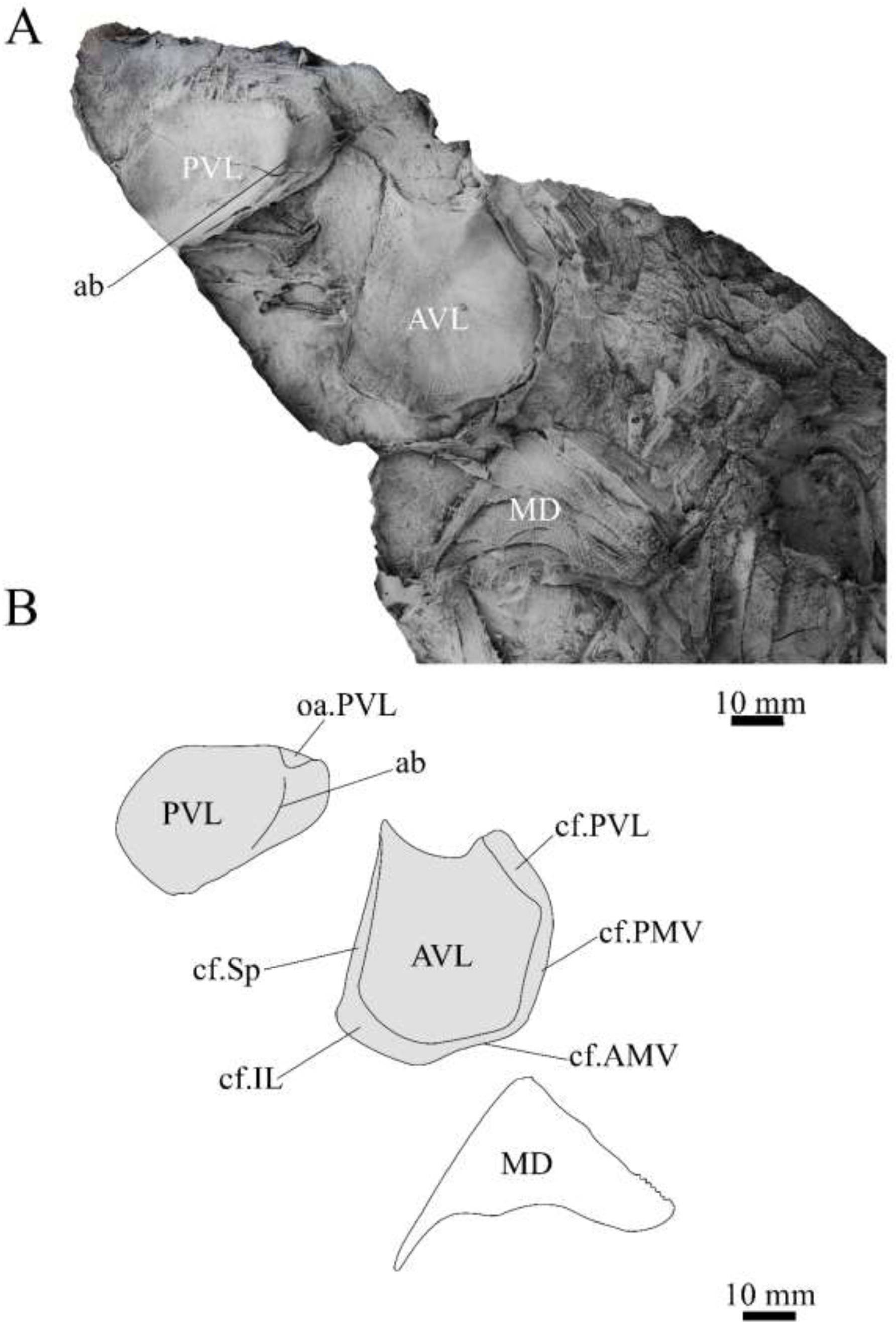
*G. howittensis* sp. nov., disarticulated trunk plates, A, NMV P254749, latex peel whitened with ammonium chloride. B, interpretive drawing of the same specimen, shaded areas indicate internal side of plate.

**Figure 10.**
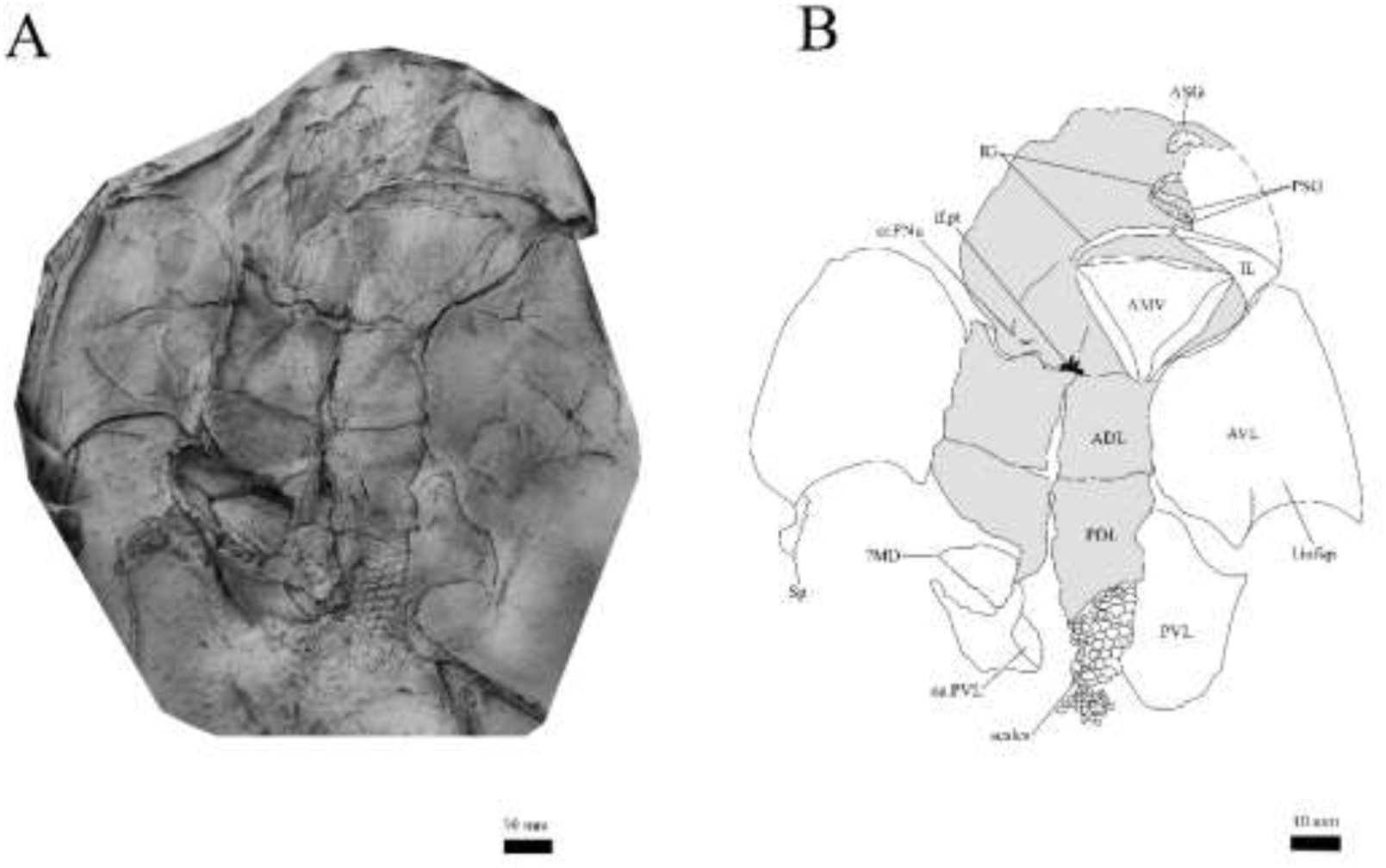
*G. howittensis* sp. nov., ventral trunk shield, ventral view. A, NMV P48884, latex peel whitened with ammonium chloride, B, interpretative drawing of the same specimen, shaded areas indicate internal side of the plate.

The parasphenoid is preserved in two specimens (Fig. 5, 7) in ventral aspect, it is a small denticulated bone, as in other groenlandaspidids, *T. elektor* (Daeschler, Frumes & Mullison, 2003) and *M. evansorum* (Ritchie, 2004). However, it is not preserved sufficiently well to provide additional anatomical detail. Visible in the holotype (Fig. 3), scattered over the ventral surface of the IG and PrO plates, are small, crenulate scales with deep surface grooves. These were possibly skin denticles covering the underside of the head.

The posterior supragnathals (PSG) are elongated, dorsoventrally flattened paired bones which almost meet on the midline, just anterior to the pineal organ. Their oral surface is entirely covered in small, densely-packed, pointed teeth that radiate from a posteromedial depression, with the largest denticles occupying the outermost margins. The posterior supragnathals of *G. howittensis* sp. nov. are almost identical in structure and position of the “supragnathals” of *T. elektor* (Daeschler, Frumes & Mullison, 2003, fig. 8) and “anterior supragnathals” of *A. doryssa* (Gess & Trinajstic 2017, fig. 2B) therefore these tooth plates are presumed homologous with the Mount Howitt species.

The infragnathal (IG) is a long and slender bone with a slight mesial curvature. The ventral surface is furrowed by a deep meckelian groove (v.gr, Fig. 2, 7) which would have housed the dorsal edge of the meckelian cartilage in life (Young *et al*., 2001). The occlusal surface of the IG, best represented by one juvenile specimen (Fig. 5), is entirely covered by short, densely packed teeth, as in phyllolepids (Long, 1984; Ritchie, 2005), thus precluding the abductor division or “non-biting portion” which characterizes the IGs of eubrachythoracid arthrodires (Stensiö, 1963). The teeth increase in size from a single posterior point suggesting tooth addition occurred posteriorly from a single ossification centre (Fig. 5).

### Trunk plates

The trunk armour consists of the same dermal plates as in other groenlandaspidids, e.g., *G. antarcticus* and *G. pennsylvanica* (Ritchie, 1975; Daeschler, Frumes & Mullison, 2003). Anterior ventral plates are absent. The posterior trunk shield exhibits a well-developed ‘annular bourrelet’, (‘b.cpd’, Goujet 1984, fig. 61B) along the posterior complex of plates (PDL, PL and PVL, Fig. 8, 9) as in other phlyctaeniids, such as *Dicksonosteus* and *Arctolepis*. The anterior dorsolateral plate (ADL) possesses a short dorsal accessory canal (acc, Fig. 1C), a feature unique to *G. howittensis* sp. nov. among members of the genus, but also present in the Early-Middle Devonian groenlandaspidid, *Mulgaspis* (Ritchie, 2004). The distinct posterior dorsolateral (PDL) is higher than long and is best preserved in NMV P48875 (H/L = 1.44, Fig. 13). The plate displays the characteristic symphysial surface for the opposite PDL (symph.s, Fig 8) and inverted V-shaped lateral line sensory canal, which are considered diagnostic for the genus (Daeschler, Frumes & Mullison, 2003; fig. 5, Janvier & Clément, 2005, fig. 8). The dorsal flexure of the lateral canal can range in angle from 96° (NMV P48875) to 105° (AMF 62437) in the examined material (the variability likely due to the angular shear of the Mount Howitt specimens e.g., Fig. 3 this article, and in *Austrophyllolepis* (Long, 1984)). The posterior lateral overlap area (oa.PL) bears a deep groove which accommodates the annular bourrelet (ab) crossing the internal surface of the posterior lateral plate (PL, Fig. 8). Much like the PDL plate the median dorsal (MD) plate is highly variable among groenlandaspidids, particularly *Groenlandaspis* (Ritchie, 1975; Janvier & Clément, 2005). The tip of the MD is usually broken in adult specimens e.g., AMF 62537 (Fig. 12) and NMV P48875 (Fig. 13) but preserved complete, however crushed, in lateral aspect in NMV P254749 (Fig. 9). In *G. howittensis* sp. nov. the plate is approximately sub-equilateral in shape (H/L = 0.65, NMV P254749, Fig. 9), its ventral margin is deeply scalloped and the ornamentation radiates from the dorsal apex of the plate developing into prominent tubercles along the caudal margin. The spinal plate (Sp) is identical to *G. antarcticus*, except for the variable presence of tiny hook-like spines on the mesial margin of the spinal plate (Fig. 1, 10, 13).

**Figure 11.**
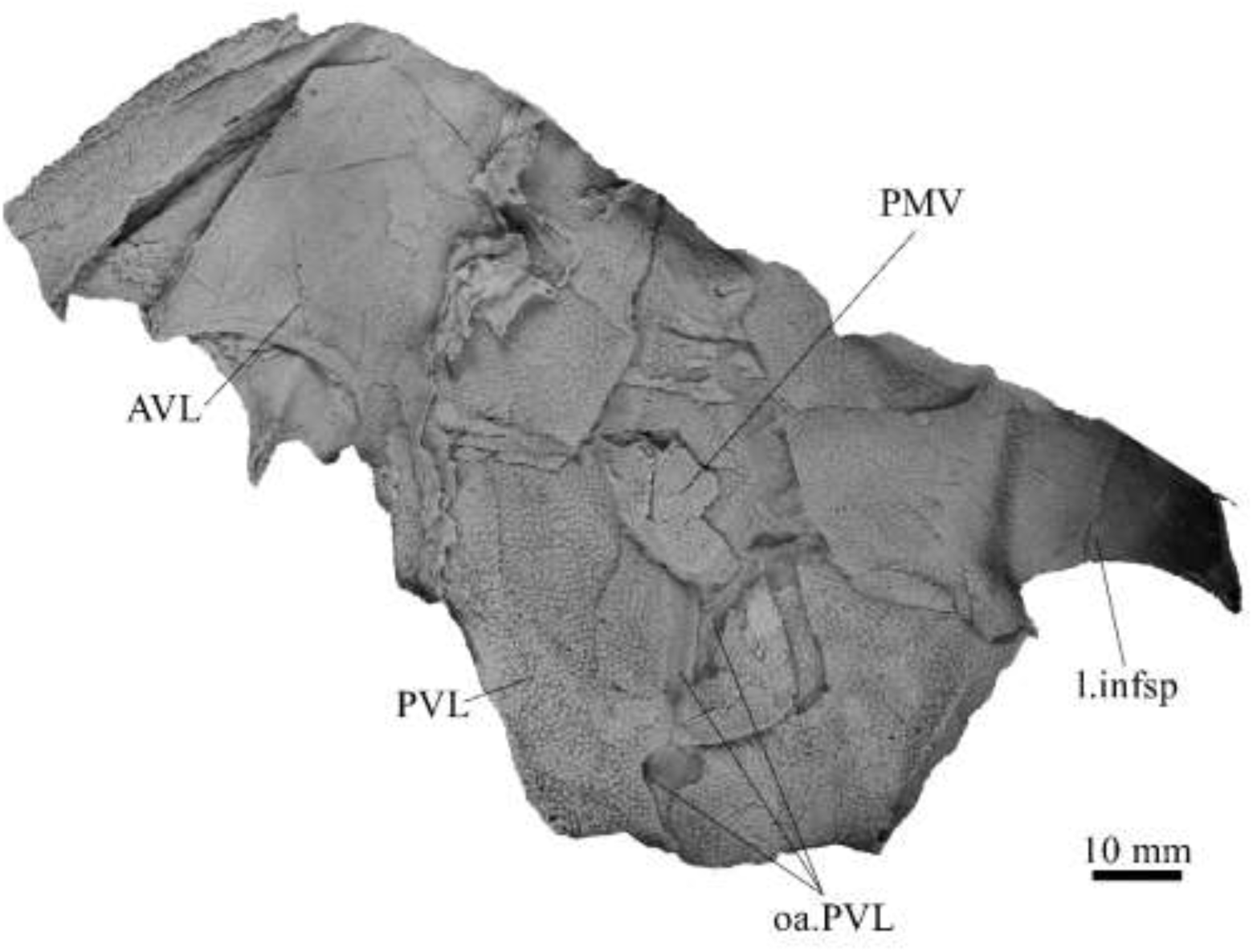
*G. howittensis* sp. nov., ventral trunk shield, ventral view, AMF 63543.

**Figure 12.**
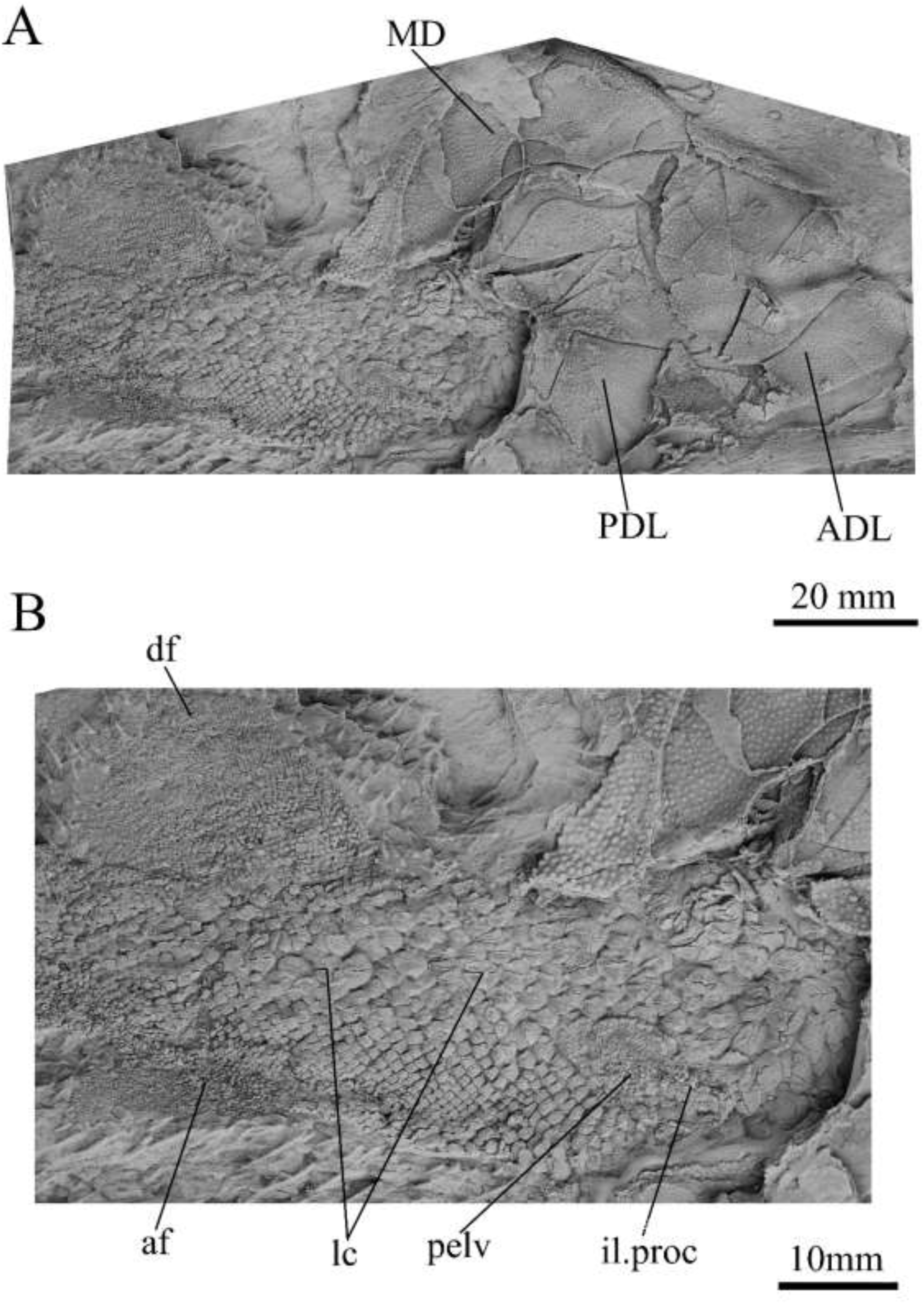
*G. howittensis* sp. nov., partial tail and lateral trunk plates, lateral view A, AMF 62537 MD, PDL, ADL and tail depicted, B, closer view of the squamation, pelvic girdle and fins of the tail. Latex peels whitened with ammonium chloride.

**Figure 13.**
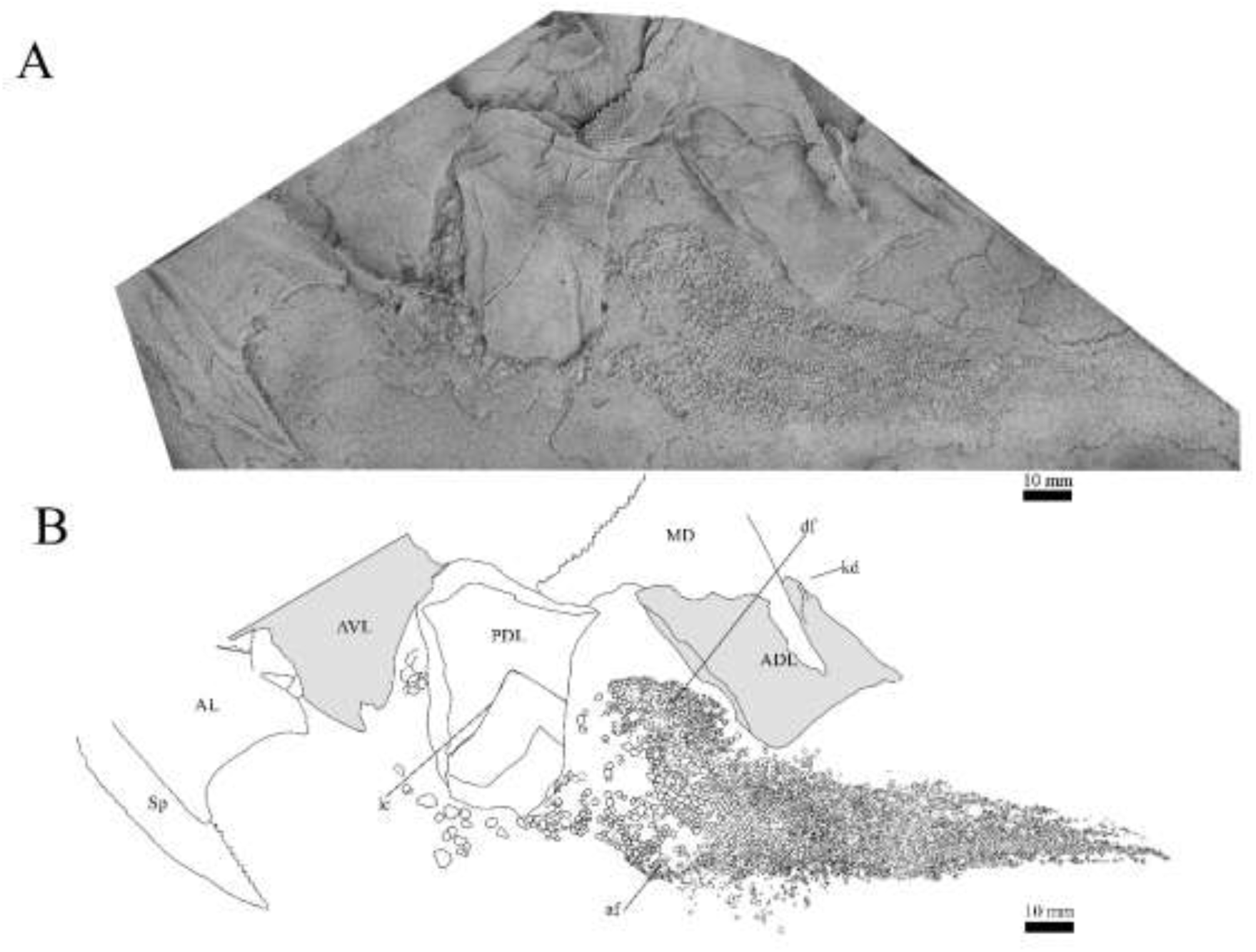
*G. howittensis* sp. nov., disaticulated trunk plates and tail in lateral view, A, NMV P48875, latex peel whitened with ammonium chloride, B, interpretive drawing of the same specimen, shaded areas indicate internal side of the plate.

The ventral surface of the trunk shield is crushed but completely preserved in the counterpart of the holotype (Fig. 1). The anterior median ventral plate (AMV) is broader than long (B/L = 1.37, NMV P48873) and similarly proportioned to other described species, *G. antarcticus* (Ritchie 1975), *G. thorezi* (Janvier & Clément, 2005), and *G. potyi* (Olive *et al*., 2015). The posterior median ventral plate (PMV) is trapezoidal and narrow (H/L = 0.53, NMV P48873).

The anterior border of the PMV and posterior border of the AMV both possess an overlap area suggesting possible midline contact of the AVL plates, though this is not confirmed in any articulated material. The posterior ventrolateral plates (PVL) exhibit a complex form of overlap areas (Fig. 11) characteristic of phlyctaeniid arthrodires (Goujet 1984).

### Pectoral Fin

The right pectoral fin is preserved as articulated dermal scales in the holotype. It is short (33mm) and broad (47mm) and covered dorsally and ventrally by small polygonal, non-overlapping scales each covered in short, rounded tubercules (Fig. 1). The pectoral fin is seldom fossilized among arthrodires, but when preserved it is typically represented by ossified endoskeletal radialia, e.g., *Incisoscutum ritchiei*, (Dennis & Miles, 1981). The pectoral fin is preserved in outline for *Amazichthys* which differs from *G. howittensis* sp. nov. in being broad and triangular in form (Jobbins *et al*., 2022).

### Post-thoracic anatomy

The tail of *G. howittensis* sp. nov. is preserved in lateral aspect in two specimens, the anterior portion in AMF 62537, (Fig. 12) and almost whole tail following the dorsal and anal fins in NMV P48875 (Fig. 13), only lacking the distal tip of the caudal fin. Both specimens are generally similarly proportioned based on comparable lengths of the MD (NMV P48875, L= 60mm and AMF 62537, L = 71mm) and thus these specimens can provide a complete restoration of the body shape and squamation for the genus (Fig. 14) and indicates a reconstructed tail length of 158mm. Based on the length of the MD (60-71mm) and tail (158mm) in these specimens summed with the average length of the skull roof (77mm) in adult specimens (NMV 48873, AMF 63542 and AMF 63535), therefore a likely overall length of *G. howittensis* sp. nov. might be between 295mm and 306mm. Not accounting for the slight downward tilt of the head which subtracts a small amount from the total length but remains unknown given the flattened nature of the fossils. Compared with other arthrodire groups where the post-thoracic region is completely known e.g., coccosteids, holonematids, phyllolepids, as well as other groenlandaspidids (*Africanaspis*) the tail of *G. howittensis* sp. nov. is relatively stout comprising roughly half the total length of the fish (Fig. 14).

**Figure 14.**
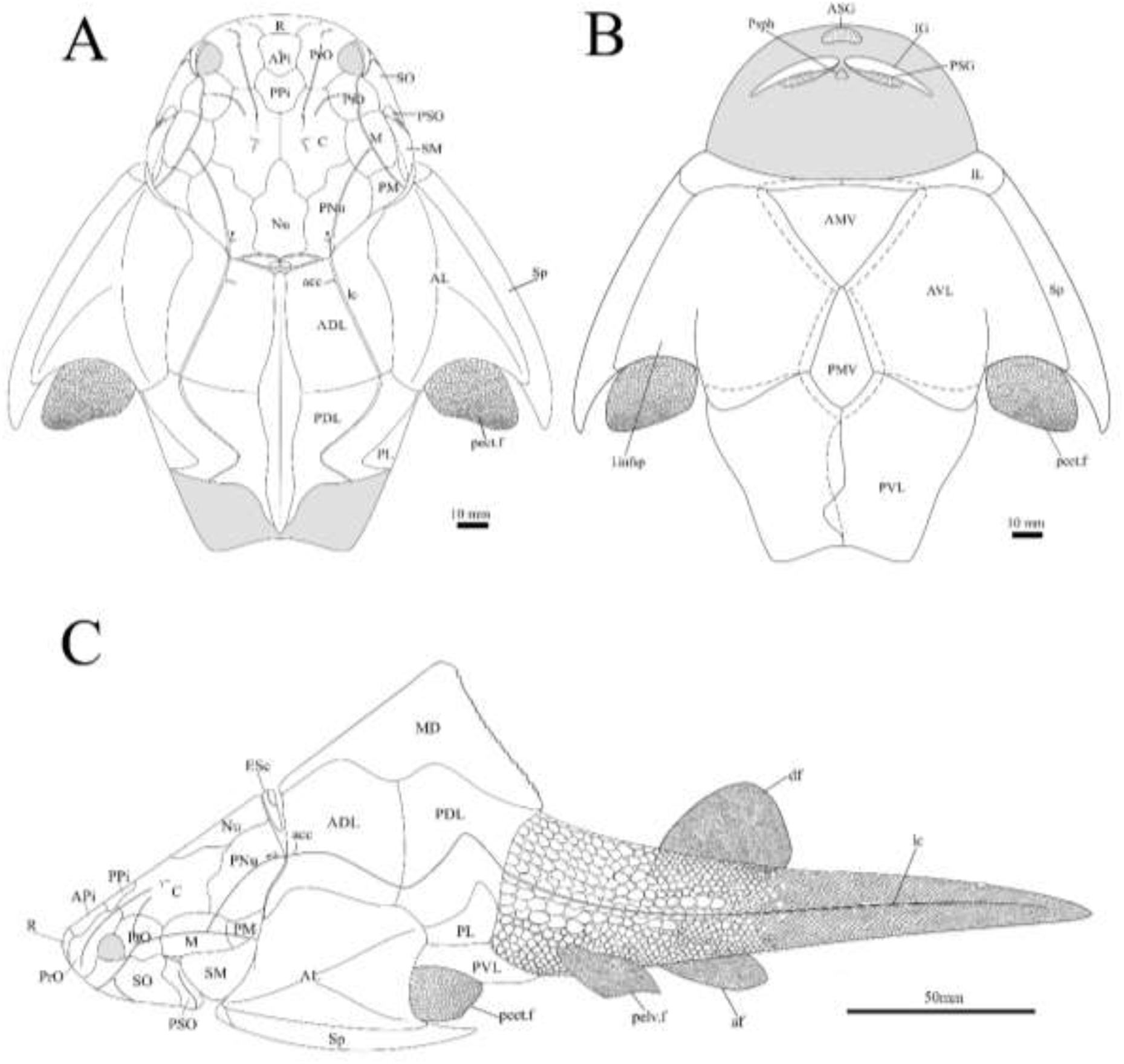
*G. howwitensis* sp. nov. reconstruction, A, dorsal view; B, ventral view; C, lateral view. dotted lines indicate overlap regions

The body scales of *G howittensis* sp. nov. display lateral and ventral variation. Burrow & Turner (1999) briefly described the lateral body scales of *G. howittensis sp. nov.* They noted the tail is covered by rhombic, non-overlapping scales 2.5-<0.1mm in length covered in and bear transverse ridge, some of these scales are deeply furrowed by the continuation of lateral canal from the PDL (Fig. 12). A postmedian “scute” (pms) can be observed toward the caudal end of NMV P48875 (Fig. 13), it is similar in morphology to the larger scales toward the base of the tail. Such “scutes” also occur in several other stem gnathostomes, e.g. *Kujdanowiaspis* and *Xuishanosteus* (Dupret *et al*., 2010; Zhu *et al*, 2022). A portion of the ventral side of the tail is preserved in one specimen, NMV P48884, wherein overlapping scales immediately posterior to the base of the PVL plates are transversely elongated and completely lack ornamentation (Fig. 10). A putative pelvic girdle is identified by a poorly-defined impression in AMF 62537 (Fig. 13). It shows a slender iliac process (il.proc) and broad basal plate (pelv) as in Gogo arthrodires, e.g., *Incisoscutum ritchiei* (Dennis & Miles, 1981) though overlying scales obscure finer anatomical detail.

### Phylogenetic Results

The results of the 50% majority rule tree (Fig. 15) include clades which are identified in the strict consensus of other analyses, e.g., Carr & Hlavin (2010) and Zhu *et al*., (2016), but are not resolved in our strict consensus due to unstable taxa. A parsimony analysis (heuristic search) of our modified data matrix returned 35234 equally parsimonious trees at 618 steps (Fig. 15). The topology of our 50% consensus analysis is broadly comparable to the strict consensus of Zhu *et al*. (2016, fig. 9) though we recover lower support values for branches concerning homostiid and dunkleosteid taxa. Additionally, the superfamily Incisoscutoidea is paraphyletic. The two Moroccan eubrachythoracids added in this analysis, *Amazichthys* and *Alienacanthus*, emerge as sister taxa nested among other aspinothoracids, in congruence with Jobbins *et al*. (2024). The node supporting the Brachythoraci is defined by two synapomorphies; a laterally expanded or trapezoidal nuchal plate (char. 105) and contact of the ADL and PL plates (char. 126). The phlyctaeniid node is supported by the following synapomorphies: midline contact of the ADLs (char. 128), an internal thickening of the posterior trunk plates (char. 129) and sigmoidal/double overlapping of the PVL plates (character 130). In the strict consensus groenlandaspidids nested among the phlyctaeniids, sister to the arctolepids (*Heintzosteus* and *Arctolepis*) with *Dicksonosteus* one node basal. The groenlandpasidid *M. evansorum* recovers most basal among groenlandaspidids, followed by *Tiaraspis* in the 50% consensus. All members of the genus *Groenlandaspis*, including *G howittensis.* sp. nov. sit crownward to other groenlandaspidids in our 50% majority rule tree except for *Africanaspis* which is recovered in a polytomy with *G. riniensis* basal one node to other species of *Groenlandaspis*. The incompletely known taxon *Elvaspis tuberculata* recovers either basal to the phlyctaeniids or basal to the brachythoracids in most parsimonious trees.

**Figure 15.**
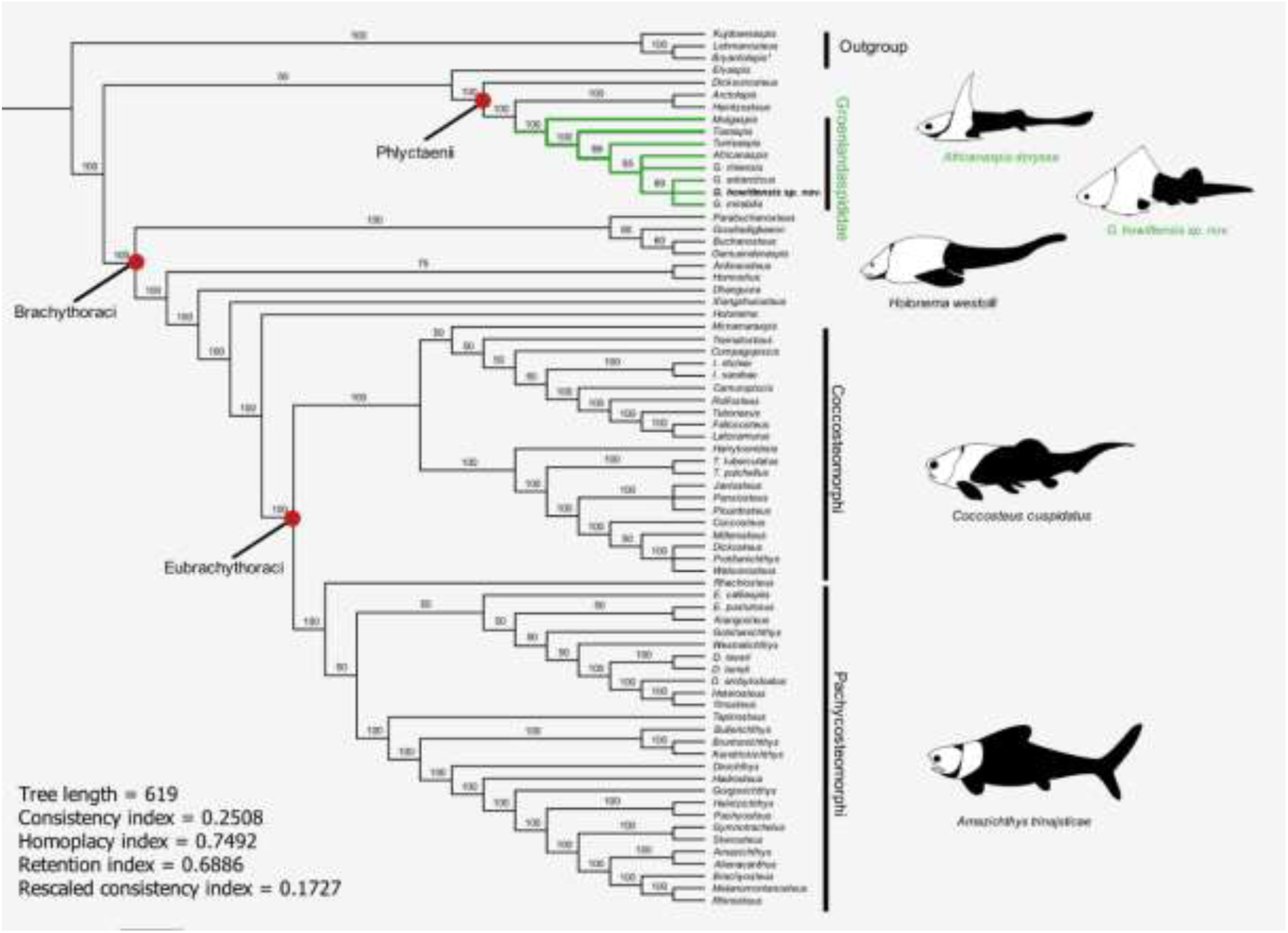
50% majority-rule consensus of 35234 equally parsimonious trees showing the phylogenetic relationships of *G. howittensis* sp. nov. and Groenlandaspididae (highlighted green) among phlyctaenioid arthrodires. Values at nodes indicate consensus frequency (thus only nodes which occur at 100% will also appear on the strict consensus). Image silhouettes are our own (*G. howittensis*) or modified from the following: *Africanaspis* doryssa, (Gess & Trinajistic 2017, fig. 3); *Holonema westolli,* (Trinajstic 1999, fig. 5C); *Coccosteus cuspidatus* (Trinajstic *et al*. 2015, fig. 16); *Amazichthys trinajsticae* (Jobbins *et al*. 2022, fig. 9).

## DISCUSSION

### Intraspecific variation

Intraspecies variation is a pervasive problem in the description of fossil organisms. Anatomically distinct specimens can appear as two taxa without the presence of intermediate forms. In some cases the geological history of a site can influence the taxanomic identity of specimens, as in, *Austrphyllolepis youngi* which was originally considered distinct from *Austrophyllolepis ritchei* (Long 1984). However, the angular shear of the deposit created distortion in the Mount Howitt specimens that was not initially recognised by Long (1984). Intraspecific variation, particularly regarding the MD plate has been recognised in other groenlandaspidids, e.g., *Turrisaspis* (Daeschler, Frumes & Mullison, 2003) and some variation is noted in the material of *G. howittensis* sp. nov.

In *G. howittensis* sp. nov. there is notable variation in the shape of the AMV plate between NMV P48874 (Fig. 1) and NMV P48884 (Fig. 10), the caudal portion of the latter being more elongate. The presence of the spinelets on the mesial margin of the spinal plate is also variable, the holotype individual lacks them NMV P48873 (Fig. 1) whereas they are clearly present on other individuals, NMV P48884 and NMV P48875 (Fig. 10, 13). Variation in the shape of the AMV has also been shown in extensive material of incisoscutid and camuropiscid arthrodires (Trinajstic & Hazelton, 2007). We equate the variance of these features to normal intraspecific variance and not substantial enough to erect an additional species though we cannot preclude the existence of two very anatomically close species of *Groenlandaspis* present in the Mount Howitt fauna.

There is also common asymmetrical variation in the path of sensory canals present on every specimen of *G. howittneiss* sp. nov. where cranial plates are preserved, e.g., on the holotype, the lateral canal (lc) of the right PNu is disjointed and in AMF 63548 (Fig. 3) the left supraorbital canal diverges briefly from its normal path. The most unusual example of this is in AMF 155378 (Fig. 8), where the PNu exhibits a second ‘aberrant canal’ (a.c) which diverges toward the post marginal canal (pmc) and does not readily compare to any before described in arthrodires. Asymmetrical variation in the growth of plates and sensory canals in arthrodires has been linked to intense environmental stresses (Trinajstic & Dennis-Bryan, 2009). This concurs with observations made of the dipnoan taxa (*Barwickia* and *Howidipterus*) of the Mount Howitt site which are thought to have recently diverged from a common ancestor driven by resource scarcity (Long & Clement 2009).

### Comparison of tooth plates with other arthrodires

Based on well-preserved examples of the tooth plates in *G. howittensis* sp. nov. it is now evident the anterior supragnathal of *Groenlandaspis* is unique among arthrodires in being a fused, medially positioned element in contrast to a generalised paired condition (Fig. 16). This specialisation has likely led to some error in the interpretation of these elements in other groenlandsaspidids. In *Turrisaspis elektor* a possible ASG is referred to as the ‘anteroventral margin of the rostral plate’ by Daeschler, Frumes & Mullison (2003). A single fused ASG was also identified by Long *et al*. (1997) in a specimen of a “juvenile *G. riniensis*”, this specimen was subsequently reassigned to *Africanaspis doryssa* by Gess & Trinajstic (2017), but not further described. Both these genera show the same unique arrangement of PSG plates as with *G. howittensis* sp. nov., supporting the likely occurrence of a fused ASG. Therefore, the presence of a dorsoventrally flattened fused ASG, should be considered a synapomorphy of the family Groenlandaspididae and present a character for analysis. In non-groenlandaspidid arthrodires, a “peg-like” fused ASG was documented for *Holonema westolli* (Miles, 1971) but subsequent newly prepared specimens form Gogo confirm it is a paired element as in other arthrodires (pers. obv.).

**Figure 16.**
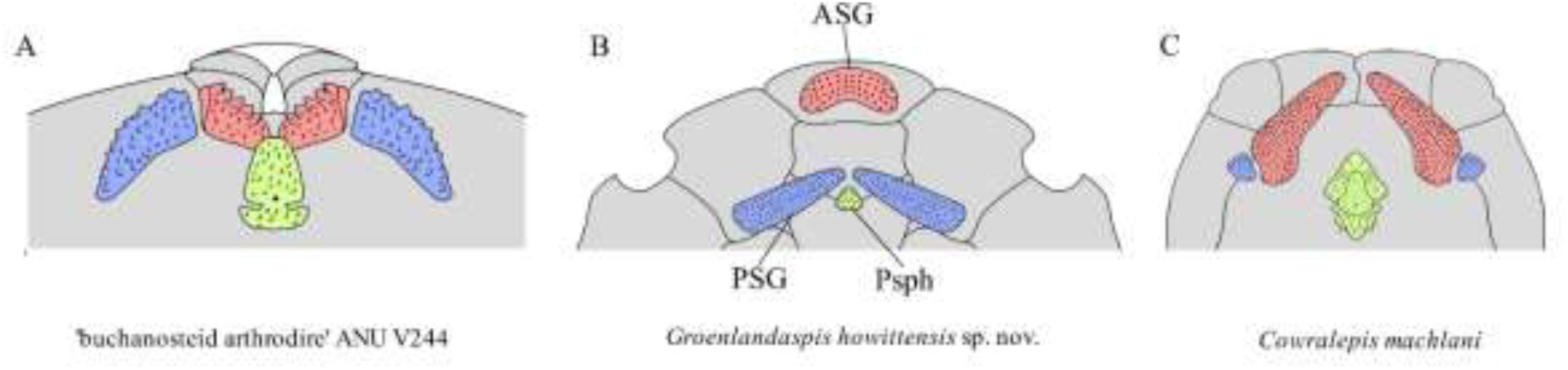
Arrangement of upper-tooth plates in basal arthrodires. Red = anterior supragnathal (ASG), blue = posterior supragnathal (PSG), green = parasphenoid (Psph). A, ‘buchanosteid arthrodire’ ANU V244, Fig. 6B. B, *Groenlandaspis howittensis* sp. nov. composite reconstruction after NMV P48773 and AMF 62534. C, *Cowralepis mclachlani* after Ritchie 2005, fig. 9F, G & 15C, D. Not to scale.

### Functional morphology and palaeoecology

The ASG bone that sits outside the main occlusion of the PSG and IG bones suggests it could be as an extra-oral element (Fig. 16). This novel adaption might have important implications for the global migration the family during the Devonian. Nonetheless, without preservation of gut contents or the remaining jaw apparatus (e.g., meckelian cartilage, palatoquadrate, hyoid arch) further inferences on the functional significance of this structure remain speculative.

The ventrally flattened body, dorsolaterally positioned eyes and ventrally positioned mouth, are consistent with bottom feeding habits and a demersal niche characteristic of basal arthrodires (Miles, 1969). A relatively stout, heavily scaled tail suggests *G. howittensis* sp. nov. was likely a weak swimmer, the short and inflexible pectoral fins likely only assisted in minor lift to keep the fish slightly above the bottom of its lacustrine habitat when it swam. The fine, tuberculate homodont dentition of this species aligns with a villiform morphotype adapted for gripping rather than crushing or puncturing prey common in extant demersal fish, e.g., groupers (*Epinephelus*, Mihalitsis & Bellwood, 2019) or siluriformes (Sado *et al*., 2020).

Alternatively, Gess & Whitefield (2020) interpreted the tooth plates of *G. riniensis* as those adapted to a durophages diet, supported by the occurrence of bivalves preserved within some juvenile specimens. A durophages habit is more likely for those groups living in marine ecosystems, whereas this contrasts with the palaeoenvironmental interpretation of the Mount Howitt site as lacustrine, with the only non-vertebrate material identified being only lycopsid plants (Long, 1983a). Moreover, the gape of *G. howittensis* sp. nov. would have been heavily limited by the narrow nuchal gap and extrascapular plates, thus, incapable of feeding on other fully-grown gnathostomes of the Mount Howitt fauna. Though the function of the peculiar tooth array cannot be further interpreted at this time, *G. howittensis* sp. nov., possibly, scoured the benthic zone for larval fishes or soft-bodied invertebrates, analogous to extant freshwater skate or catfish.

### Systematic implications

The material of *G. howittensis* sp. nov. is the most completely known example of any groenlandaspidid described and is the first member of the cosmopolitan genus *Groenlandaspis* to be formally described from Australia.

Extrascapular plates have previously been considered a specialisation of the brachythoracids (Miles, 1973; Dennis & Miles, 1979; Gardiner & Miles 1990), however, these elements have since been recognised in multiple genera of actinolepidids, e.g. *Sigaspis*, *Aleosteus,* and *Erikaspis* (Goujet, 1973; Johnson *et al*., 2000; Dupret *et al*., 2007), and now the phlyctaeniid, *Groenlandaspis*, supports extrascapular elements as being plesiomorphic for arthrodires and so subsequently lost in numerous later groups. The occurrence of these plate however presents a challenging character for analysis as they greatly affected by preservation bias. Of eight articulated specimens examined for this study only three occurrences of extrascapular plates were identified in the *G. howittensis* sp. nov. material.

King, Hu & Long (2018) reviewed the presence of possible electro sensory organs in Paleozoic gnathostomes. They noted the potential phylogenetic significance of cutaneous sensory pits (char. 126) in arthrodires. This feature is generally restricted to buchanosteids, coccostemorphs along with *Eastmanosteus* in our analysis, is variably present among *G. howittensis* sp. nov. individuals. The cheek plates for other groenlandaspidids are poorly known but these elements as described for *G. riniensis* (Long *et al*. 1997, fig. 5H) and *Africanaspis* (Gess & Trinajstic 2017, fig. 5 B, D) show no evidence of sensory pits.

The infraorder Phlyctaenii Miles 1973 is often considered as a grade group by several workers (e.g., Dennis & Miles, 1979; Gardiner & Miles, 1990; 1994 and Zhu *et al*., 2016). Our hypothesis of arthrodire phylogenetic relationships reflects that of Goujet (1984) and Dupret (2004) in supporting a monophyletic relationship of the phlyctaeniid families, Groenlandaspididae, Arctaspididae and Arctolepidae united by the specialisations: medial contact of the ADL plates, followed by contact of the PDL plates in groenlandaspidids (char. 126) and sigmoidal/ double-overlappingcontact of the PVL plates (char. 129). Although Goujet (1984) also proposed an anterior narrowing of the median dorsal plate as a synapomorphy, we consider this character functionally correlated with the medial contact of the ADLs and so it is not considered as a separate character in this analysis. Another major arthrodire family considered among the Phlyctaenii are the Phlyctaeniidae, Fowler 1947, (e.g., *Phlyctaenius* and *Pagaeaaspis*), they lack the unusual overlap pattern of the PVL plates (Young, 1983) and it is unclear if they possess a developed annular bourrelet as in *Arctolepis*, *Dicksonosteus* and *Groenlandaspis*. We propose these forms require further investigation of their phylogenetic relationships, as they are generally conceded as a grade group by other workers positioned basal to the rest of Phlyctaenioidei (Goujet, 1984; Dupret *et al*., 2017).

Our 50% consensus analysis fails to support the monophyly of the genus *Groenlandapsis* and we do not identify any unique specialisations shared between currently described members of the genus. Though we have provided an amended diagnosis we note that multiple species of *Groenlandaspis* await further description, namely, *G. disjectus* from the Kiltorcan Formation, Ireland (Ritchie, 1974), *Groenlandaspis sp*. from the Adolphspoort Formation, South Africa (Anderson *et al*., 1999), *Groenlandaspis sp*. from Canowindra, Australia and an abundance of fragmentary material from multiple other sites in Australia (Young, 1993). As such, our diagnosis for *Groenlandaspis* should be considered tentative. Furthermore, revision of the type species *G. mirabilis* is also necessary as some bones remain misidentified, e.g., the “AMV” and “AVL” only depicted by drawings in, Heintz, 1932, Fig. 12, differ strongly in shape from any known arthrodires and are likely erroneously labelled PVL plates. A full taxonomic review of *Groenlandaspis* is required to complete a definition of the genus and further probe its phylogenetic relationships.

Our analysis does not support a grouping of the three ‘high-spired’ genera with tall MD plates, *Tiaraspis*, *Turrisaspis* and *Africanaspis* as previously proposed (Olive *et al*., 2015). Gess & Trinajstic (2017) discussed similarities of these taxa, primarily the presence of a dorsolateral ridge, dual pineal elements, and the foreshortened trunk armour. Dual pineal elements (char. 122) are now properly described in *Groenlandaspis* and is likely a synapomorphy uniting a clade of derived groenlandaspidids, with a single element exhibited by *Arctolepis* and *Mulgaspis* being the plesiomorphic state. A dorsolateral ridge (char. 126) commonly reported among phlyctaeniid taxa, e.g. *Denisonosteus* (Young & Gorter, 1981) and *Phlyctaenius* (Young, 1983), yet lost in *Mulgaspis* and some species of *Groenlandaspis* is also supported by our analysis as plesiomorphic (Long, 1995). Lastly, compared to *Groenlandaspis*, the trunk armour of *Turrisaspis* and *Africanaspis* and to a lesser extent *Tiaraspis* are foreshortened in proportions, particularly in the median dorsal plate (Long *et al*., 1997; Daeschler, Frumes & Mullison, 2003). Though similarly foreshortening is present in some *Groenlandaspis* species, as in the ADL and PDL of *G. riniensis* (Long *et al*., 1997, fig. 7A, B) and the MD of *G. seni* (Janvier & Ritchie, 1977, fig. 1B, C). Signifying this morphology requires further investigation to quantify the effect of bone proportions on the phylogeny of groenlandaspidids. Also significant for the evolution of groenlandaspidids is the inflexion of the PDL sensory canal (Long, 1995). It is wide in Early-Middle Devonian groenlandaspidids, *Mulgaspis*, *Tiaraspis* and *Boomeraspis* (Long, 1995; Ritchie, 2004) and sharply flexed in certain Middle-Late Devonian forms, like *Groenlandaspis*, *Turrisaspis*, and *Africanaspis* (Daeschler, Frumes & Mullison, 2003). A wide flexion better compares with the straight canal in exhibited by many phlyctaeniids, e.g. *Dicksonosteus* (Goujet, 1984), suggesting this to be the plesiomorphic state.

Alternative hypotheses regarding the phylogenetic relationships of *Groenlandaspis* includes a grouping with *Holonema* and *Arctolepis* (Denison, 1978; 1984; Young & Gorter, 1981) in the family Holonematidae chiefly based on the putative fusion of the postnasal bones with the rostral plate. Though a compound rostral and postnasal bone is supported in the Gogo material for *Holonema* (Miles, 1971), Goujet (1984) found no evidence of this in *Arctolepis* and nor do we for *Groenlandaspis*. Our strict consensus places *Holonema westolli* within Brachythoraci, further crownward than the buchanosteids, and supports Miles’ (1971) interpretation of the genus as an early diverging brachythoracid.

## CONCLUSION

*G. howittensis* sp. nov. provides us with rare insight into the morphology of the post-trunk skeleton, fins and dental morphology for arthrodires. The exceptional preservation of the Mount Howitt specimens reveals undescribed details of the tooth plates for groenlandaspidids, highlighting a uniquely specialised condition where the ASG is fused and positioned anterior to the remainder of the tooth arcade. *G. howittensis* sp. nov. is a unique example of extreme dental specialisation and evolutionary experimentation in stem jawed vertebrates nearing the origin of teeth. The phylogenetic relationships of the Groenlandaspididae are presented for the first time in a computer-driven phylogenetic analysis.

## ACKNOLWEDGMENTS

We are grateful to Dr Matthew McCurry, of the Australian Museum for graciously making latex peels of many specimens in their collection. We thank Tim Ziegler for providing access to the palaeontological collections of the Melbourne Museum and for his assistance in locating specimens. We thank Shona Ritchie and the Canowindra Age of Fishes Museum for access to the notes and casts of specimens made by the Dr Alex Ritchie. This research was funded by a donation of funds to the research account of one of us, JAL, by businessman John Clema of New Mexico, USA.

## Institutional Abbreviations

NMV: Museum of Victoria, Melbourne, Australia

AMF: Australian Museum, Sydney, Australia

ANU: Australian National University, Canberra, Australia

## Anatomical Abbreviations

ab: annular bourrelet

a.c: aberrant canal

acc: accessory canal

ADL: anterior dorsolateral plate

af: anal fin

AL: anterior lateral plate

AMV: anterior median ventral plate

APi: anterior pineal plate

ASG: anterior supragnathal

AVL: anterior ventrolateral plate

C: central plate

cf.ADL: contact face for the anterior dorsolateral plate

cf.AMV: contact face for the anterior median ventral plate

cf,MD: contact face for the median dorsal plate

cf.PDL: contact face for the posterior dorsolateral plate

cf.IL: contact face for the interolateral plate

cf.PMV: contact face for the posterior median ventral plate

cf.PVL: contact face for the posterior ventrolateral plate

cf.Sp: contact face for the spinal plate

csc: central sensory canal

cr.PNu: paranuchal crista

cuso: cutaneous sensory organ

df: dorsal fin

end.d: endolymphatic duct

Esc: extrascapular plates

Esc.c: extracapsular plate canal

if.pt: infranuchal pit

IG: infragnathal

IL: interolateral plate

il.proc: iliac process of the pelvic gridle

ioc: infraorbital canal

kd: articular condyle

lc: lateral canal

l.infsp: infraspinal lamina

MD: median dorsal plate

mpl: median pit line

Nu: nuchal plate

oa.AVL: overlap area for the anterior ventrolateral plate

oa,C: overlap area for the central plate

oa.IL: overlap area for interolateral plate

oa.M: overlap area for the marginal plate

oa.N: overlap area for the nuchal plate

oa.PL: overlap area for the posterior lateral

oa.PVL: overlap area for the posterior ventrolateral plate

occ: occipital cross commissure

orb: orbit

pap: para-articular process

PDL: posterior dorsolateral plate

pdl: posterior descending lamina

pect.f: pectoral fin

pelv: basal plate of the pelvic girdle

pelv.f: pelvic fin

PL: posterior lateral plate

PPi: posterior pineal plate

ppl: posterior pit line

ppt: pineal pit

psoc: post suborbital canal

PM: post marginal plate

pmc: postmarginal canal

pms: post median scute

PMV: posterior median ventral plate

PNu: paranuchal plate

PrO: preorbital plate

PSG: posterior supragnathal

PSO: post suborbital plate

Psph: parasphenoid

PtO: postorbital plate

PVL: posterior ventrolateral plate

R: rostral plate

SM: submarginal

SO: suborbital

soc: supraorbital canal

sorc: supraoral canal

Sp: spinal plate

suo.v: supra orbital vault

symph.s: symphysial surface

v.gr: ventral groove.

PLACODERMI McCoy, 1848

ARTHRODIRA Woodward, 1891

PHLYCTAENIOIDEI Miles, 1973

PHLYCTAENII Miles, 1973

GROENLANDASPIDIDAE Obruchev, 1964

*GROENLANDASPIS* Heintz, 1932

