## Supplementary material for "Unique dental arrangement in a new species of *Groenlandaspis* (Placodermi, Arthrodire) from the Middle Devonian of Mount Howitt, Victoria, Australia": SImatrixCorrections.docx

Modifications from Zhu *et al.* (2016) phylogenetic matrix

*Dicksonosteus arcticus*

( 1 ) 0 > 1

(51) 0 > 1

(63 ) ? > 1

(77) 1 > 0

*Groenlandaspis antarcticus*

(24) 1 > 0

(26) 0 > -

(30) 0 > 1

(51) ? > 1

(53) ? > 0

(77) 1 > 0

*Turrisaspis elektor*

(13) 0 > 1

(20) 0 > 1

(26) 0 > -

(50) ? > 0

(53) ? > 0

(77) 1 > 0

*Elvaspis tuberculata*

(51) 0 -> 1

*Kujdowniaspis podolica*

(11) 0 > 2

(31) 1 > 0

Zhu YA, Li Q, Lu J, Chen Y, Wang J, Gai Z, Zhao W, Wei G, Yu Y, Ahlberg PE, Zhu M. 2022. The oldest complete jawed vertebrates from the early Silurian of China. *Nature* 609(7929):954-958. [DOI: 10.1038/s41586-022-05136-8](https://doi.org/10.1038/s41586-022-05136-8)
